# H4K16ac activates the transcription of transposable elements and contributes to their cis-regulatory function

**DOI:** 10.1101/2022.04.29.488986

**Authors:** Debosree Pal, Manthan Patel, Fanny Boulet, Jayakumar Sundarraj, Olivia A Grant, Miguel R. Branco, Srinjan Basu, Silvia Santos, Nicolae Radu Zabet, Paola Scaffidi, Madapura M Pradeepa

## Abstract

Mammalian genomes harbour a large number of transposable elements (TEs) and their remnants. Many epigenetic repression mechanisms are known to silence TE transcription. However, TEs are upregulated during early development, neuronal lineage, and cancers, although the epigenetic factors contributing to the transcription of TEs have yet to be fully elucidated. Here we demonstrated that the male-specific lethal (MSL) complex mediated acetylation of histone H4 lysine 16 (H4K16ac) activates transcription of long interspersed nuclear elements (LINE1, L1) and long terminal repeats (LTRs). Furthermore, we show that the H4K16ac marked L1 and LTR subfamilies function as enhancers and are enriched with chromatin features associated with active enhancers and looping factors. L1 and LTRs enriched with histone acetylations are bound by chromatin looping factors and these regions loop with genes. CRISPR-based epigenetic perturbation and genetic deletion of L1s reveal that H4K16ac marked L1s and LTRs regulate the expression of genes in cis. Overall, TEs enriched with H4K16ac contribute to the cis-regulatory landscape of a significant portion of the mammalian genome by maintaining an active chromatin landscape at TEs.

**One Sentence Summary:** H4K16ac activates LINE1 and ERV/LTR transcription and rewires the cis-regulatory landscape of a significant portion of the mammalian genome by increasing the transcriptional activity at TEs.

## Introduction

Dysregulation of TEs and their insertions into gene exons are usually disruptive and are implicated in cancer and neurological disorders (Burns, 2017; Hancks and Kazazian, 2016). When inserted into noncoding DNA, including introns, they can affect the host gene expression in *cis* or *trans*. Most TEs are incapable of transposing due to acquired mutations and epigenetic and post-transcriptional silencing mechanisms (reviewed in)(Almeida et al., 2022; Molaro and Malik, 2016). Transcription of TEs is repressed by DNA methylation, H3K9me3, TRIM28/KRAB-ZFPs, and HUSH complex (Bulut-Karslioglu et al., 2014; Karimi et al., 2011; Robbez-Masson et al., 2018; Rowe et al., 2010; Walsh et al., 1998). Apart from these repressive mechanisms, several pluripotency-associated transcription factors, SP1/SP3, LBP9, DUX4, DUX, GATA2 and YY1, are enriched at ERV/LTRs, and SOX11, RUNX3, YY1 are enriched at the 5’ UTRs (promoters) of LINE1 (L1) (reviewed in) (Hermant and Torres-Padilla, 2021). Interestingly, most of the species-specific DNase hypersensitive sites (accessible chromatin) are occupied by remnants of TEs (Jacques et al., 2013; Vierstra et al., 2014), suggesting that TEs have co-opted as tissue and species-specific gene regulatory elements. TEs are transiently upregulated during the early development (Jachowicz et al., 2017), in the neuronal lineage (Upton et al., 2015) and cancer (Hancks and Kazazian, 2016). The endogenous retrovirus (ERV) superfamily of long-terminal repeats (LTR) and short interspersed nuclear element (SINE/*Alu*) often exhibit chromatin features associated with active cis-acting regulatory elements (CREs) (Fueyo et al., 2022; Sundaram and Wysocka, 2020; Todd, Christopher D, Taylor and Branco, 2019) and function either as enhancers to regulate genes in cis or act as alternate promoters (Fueyo et al., 2022). 5’ UTR of long interspersed nuclear element 1 (LINE-1, L1) repeats are also bound by tissue-specific transcription factors (TFs) and can function as nuclear noncoding RNAs (Jachowicz et al., 2017; Percharde et al., 2018); still, it is unclear whether they can act as CREs. Although TEs are suggested to contribute to nearly one-quarter of the regulatory epigenome (Chuong et al., 2017; Hermant and Torres-Padilla, 2021; Jachowicz et al., 2017; Schmidt et al., 2012). The chromatin-based mechanisms contributing to regulatory activity in this vast number of TEs are unclear.

Chromatin features such as a combination of H3K4me1 and H3K27ac, bidirectional transcription of enhancer RNAs (eRNAs) and accessible chromatin (e.g., using ATAC-seq) are widely used to predict enhancer activity, including for TE-derived enhancers (Andersson et al., 2014; Buenrostro et al., 2013; Creyghton et al., 2010; Deniz et al., 2020; Todd, Christopher D, Taylor and Branco, 2019). Yet the level of H3K27ac does not correlate with or is dispensable for enhancer activity, suggesting that other uncharacterised chromatin features could contribute to regulatory activity (Kheradpour et al., 2013; Taylor et al., 2013; Wang et al., 2022). H4K16ac and H3K122ac are particularly interesting among many histone acetylations as they directly alter chromatin structure and increase transcription *in vitro* (Shogren-Knaak et al., 2006; Tropberger et al., 2013). H4K16ac and H3K122ac are enriched at enhancers, and they identify new repertoires of active enhancers that lack detectable H3K27ac (Pradeepa et al., 2016; Taylor et al., 2013). However, it is challenging to decipher the causal role of specific histone acetylations, as many acetylations, including H3K27ac, are catalysed by multiple lysine acetyltransferases (KATs), and KATs also have a broad substrate specificity. H4K16ac is an exception, as it is catalysed explicitly by KAT8 when associated with a male-specific lethal (MSL) complex.

Nevertheless, when KAT8 is associated with non-specific lethal (NSL), it catalyses H4K5ac, H4K8ac and H4K12ac (Chatterjee et al., 2016; Chelmicki et al., 2014; Radzisheuskaya et al., 2021; Ravens et al., 2014). In mouse embryonic stem cells (mESCs), KAT8 and H4K16ac mark active enhancers and promoters of genes that maintain the identity of the mESCs (Li et al., 2012; Taylor et al., 2013). Loss of function mutations in *KAT8* or *MSL3* leads to reduced H4K16ac levels and is known to cause neurodevelopmental disorders (Basilicata et al., 2018; Li et al., 2020). However, the mechanism through which MSL/KAT8-mediated H4K16ac contributes to genome regulation during normal development is less clear, especially in the human genome.

Here, we show that H4K16ac is enriched at L1 and LTRs and depleted at gene promoters, and H4K16ac regulates transcription across the L1 and ERV/LTR superfamily of TEs. TEs marked with acetylations loop with the genes and regulate their expression. CRISPR interference and genetic deletion of H4K16ac marked (H4K16ac+) L1s leads to the downregulation of genes in cis, demonstrating that H4K16ac+ TEs function as enhancers. Furthermore, depletion of H4K16ac is sufficient for genome-wide downregulation of L1 and LTRs and genes linked with these TEs, confirming the significance of H4K16ac-mediated activation of TEs in rewiring the regulatory landscape of a significant fraction of the mammalian genome.

## Results

We aimed to investigate the role of MSL-mediated H4K16ac in human genome regulation. We performed two to three replicates of Cleavage Under Targets and Tagmentation (CUT&Tag) (Kaya-okur et al., 2019) in human embryonic stem cells (H9-hESCs) for histone modifications that associate with active regulatory elements (H3K27ac, H3K122ac, H4K12ac, H4K16ac, H3K4me1 and H3K4me3), polycomb repressed domains (H3K27me3) and heterochromatin (H3K9me3) (fig. S1A). We evaluated the overall data quality, and similarity among our CUT&Tag replicates (fig. S1A). We generated peaks by merging the replicates and used reproducible peaks in at least two replicates to validate our findings (fig. S1, S2, table S2 and S3). To prevent the same reads from mapping to multiple regions in the repeat elements, uniquely mapped CUT&Tag sequencing reads were used for all analyses. We have also used total mapped CUT&Tag and RNAseq reads for mapping to L1 subfamilies to avoid bias.

### H4K16ac and H3K122ac are enriched at LINE1, ERV/LTR and SINE elements

Chromatin-state discovery and genome annotation analysis (ChromHMM) of CUT&Tag peaks revealed the expected enrichment of H3K4me1, H3K4me3, H3K27ac and H4K12ac at chromatin features associated with active transcription, including active promoters and enhancers. Intriguingly, H4K16ac and H3K122ac, but not H3K27ac or H4K12ac, were enriched at heterochromatin, insulator, and transcription elongation states (Fig. 1A). Furthermore, we found specific enrichment of H4K16ac and H3K122ac at the 5’ UTR of full-length L1s and ERV/LTR elements compared to gene promoters (Fig. 1B-E). H4K16ac is also detected at gene bodies, consistent with the previous findings showing its role in the transcription elongation (Larschan et al., 2011) (Fig. 1B). Interestingly, however, H4K16ac shows a very low level of enrichment at the gene promoters (Figs. 1B, D and E, table S3), which is similar to the recent ChIPseq data in the human cell lines (Radzisheuskaya et al., 2021). H3K9me3 level is lower at H4K16ac peaks that overlap with L1 5’ UTRs, and the H4K16ac level is lower at L1s with H3K9me3 peaks (fig. S1C). Although H4K16ac and H3K9me3 are enriched at L1s (Fig. 1B), they are anti-correlated at L1 subfamilies.

**Fig. 1:**
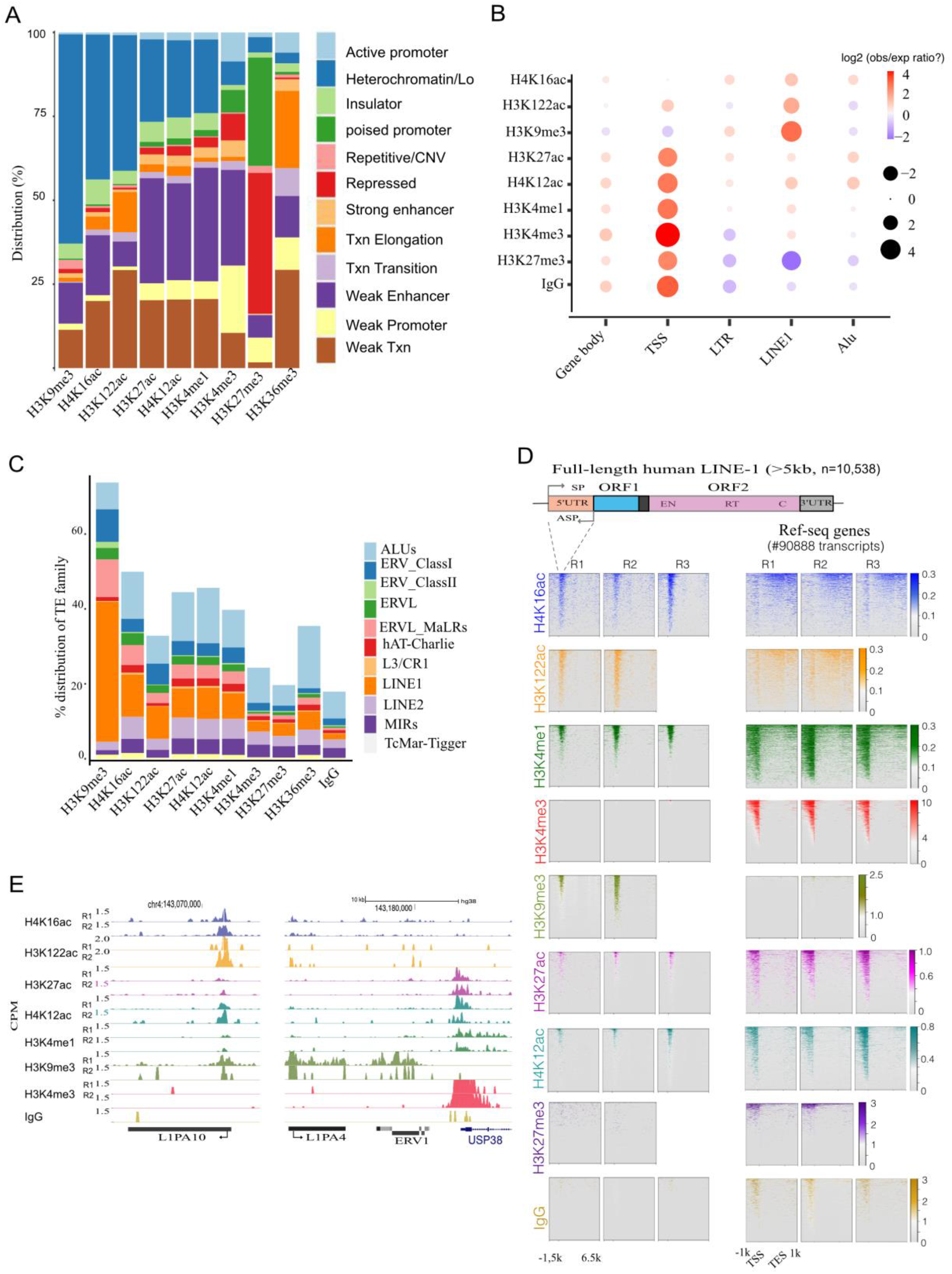
H4K16ac and H3K122ac are enriched at 5’ UTR of L1 and ERV/LTRs in hESCs A. **A**. Bar chart showing the percentage distribution (Y-axis) of histone PTMs CUT&Tag peaks across ChromHMM chromatin features. **B**. Dot plot showing the ratio (observed/expected) of enrichment of CUT&Tag peaks across gene transcription start sites (TSS), TE families (L1, ERV/LTRs, SINE/*Alu*) and gene-body. The size of the circle represents the log2 value for the ratio, and the colour range represents the gradient for the enrichment. **C**. Percentage distribution of repeat elements for CUT&Tag peaks. **D**. Illustration showing the structure of human L1s (above). Heatmap displaying the histone modifications (all replicates) CUT&Tag signal (counts per million, CPM) at full-length L1 (>5kb, left) and NCBI Ref-seq genes (right). **E**. UCSC genome-browser tracks showing signal density (CPM) of histone modifications (individual replicates) at representative L1 subfamily, L1PA10 (left), and L1PA4, ERV1 and USP38 gene (right).

Reanalysis of public ChIPseq datasets showed enrichment of H4K16ac at the 5’ UTRs of L1s in the human brain tissue (fig. S3A) (Nativio et al., 2018). H4K16ac is also enriched at the 5’ UTR of L1s in neuroblastoma (SH-SY5Y), erythroleukaemia (K562) and transformed dermal fibroblasts (TDFs) cell lines and mESCs (fig. S3B and C). This analysis suggests that H4K16ac enrichment at TEs is not unique to hESCs but is conserved in cancer cells, human brain tissue, and mice. Although H3K27ac and H4K12ac are detected at L1 5’ UTRs, they are enriched at a much higher level at the promoters of genes than L1s (fig. 1B). Interestingly, along with H4K16ac and H3K122ac, L1 5’ UTRs are also enriched with H3K4me1 and are depleted of H3K4me3 (Fig. 1C and D), suggesting that these elements could function as cis-regulatory elements (see in Fig. 3).

### H4K16ac+ L1 5’ UTRs are enriched with enhancer chromatin features

LTR subfamilies function as enhancers to regulate genes in a tissue-specific manner in humans and mice (Deniz et al., 2020; Todd et al., 2019). LTR5- and LTR7-related subfamilies function as enhancers in hESCs (Chuong et al., 2017; Fuentes et al., 2018; Pontis et al., 2019). However, it is unknown whether L1 elements can act as enhancers to regulate genes in cis. Here we found H4K16ac is particularly enriched at the 5’ UTR of full-length L1 subfamilies and correlates with chromatin features associated with active enhancers such as H3K27ac, H3K4me1, BRD4 and ATACseq signal (Fig. 2A and D) but are devoid of promoter mark H3K4me3 in hESCs (Fig. 1D, E and 2A, 2D).

**Fig. 2.**
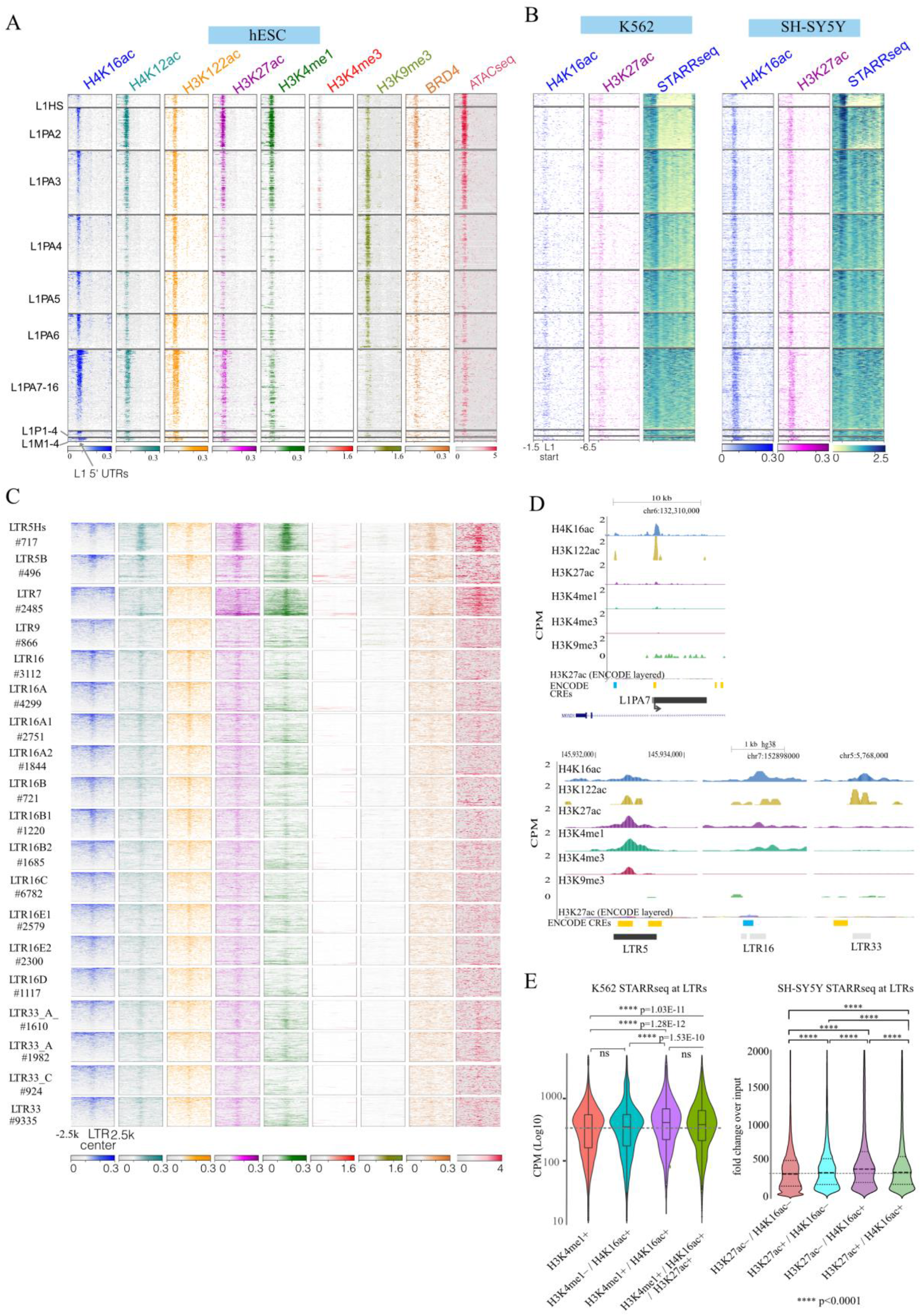
H4K16ac+ TEs are enriched with chromatin features associated with enhancer activity. **A**. Heatmap of CUT&Tag signals for histone modifications and BRD4 normalised to IgG and ATACseq signal at TE sub-families; –1.5kb to +6.5kb from the full-length L1 start sites (>5kb). **B**. Heatmap shows H4K16ac and H3K27ac CUT&Tag signals and STARRseq signals in K562 and SH-SY5Y cells. **C**. Like A, but for ±2.5kb around ERV/LTR center for subfamilies of LTR5, 7, 9, 16, 33 and MER21C. Data for the rest of the LTR and *Alu* sub-families in fig. S5). **D**. Genome browser tracks showing average (n=2 or more) CPM for two replicates of H4K16ac, H3K122ac, H3K27ac, H3K4me1 and H3K4me3 CUT&Tag data from hESCs. RepeatMasker tracks showing L1 (L1PA7, top panel), LTR5, LTR16 and LTR33 (bottom panel) and ENCODE layered H3K27ac and cis-regulatory elements (CRE) are shown below each panel. **E**. Violin plots showing STARRseq signal from K562 cells across LTRs that intersect with-H3K4me1 peaks; H4K16ac but not H3K4me1 peaks; H3K4me1 and H4K16ac peaks; and H3K4me1 along with both H3K27ac and H4K16ac peaks (Dunn’s test with Bonferroni correction).

Interestingly not all L1 subfamilies are enriched with active enhancer features at the same level. The young L1s (L1HS, L1PA2 and L1 PA3) are enriched with active enhancer features, including H4K16ac. Young L1s are known to be transcriptionally active. Despite older L1s being transcriptionally inactive, 5’ UTRs of these L1 subfamilies (L1PA7-L1PA16) are enriched with H4K16ac along with other active enhancer features but are depleted of H3K9me3 (Fig. 2A), suggesting that these 5’ UTR of older full-length L1s have been co-opted to function as functional regulatory elements.

Analysis of genome-wide enhancer activity data (STARRseq) generated by ENCODE (Lee et al., 2020) from neuroblastoma (SH-SY5Y) and erythroleukemia (K562) cell lines, showed enhancer activity specifically at the 5’ UTR of the L1s (Fig. 2B) in a cell type-specific manner. The presence of active enhancer chromatin features (Fig. 2A), together with the ability of L1 5’ UTR to drive transcription of the minimal promoter in an in vitro enhancer reporter assay (Fig. 2B), further confirmed that 5’ UTRs of full-length L1s could function as transcriptional enhancers.

### LTRs marked with H4K16ac could function as active enhancers in hESCs

Apart from LTR5 and LTR7 elements that show clear enrichment of active enhancer chromatin features, our data show that some of the subfamilies of LTR16 and LTR33 may also serve as enhancers as they are enriched with H4K16ac and other active enhancer chromatin features (Fig. 2C, D). Interestingly, STARRseq data from K562 and SH-SY5Ycells revealed H4K16ac+ LTRs show significantly higher enhancer activity than LTRs that overlap with only H3K4me1 or H3K27ac peaks (Fig. 2E and fig. S4A). These results further support that H4K16ac+ LTRs function as enhancers. The rest of the LTR and *Alu* families are not likely to act as enhancers in hESCs, as they lack known enhancer chromatin features (fig. S4B).

### L1/LTRs marked with H4K16ac are enriched with chromatin looping factors

We aimed to identify TFs bound at H4K16ac+ TEs along with H3K27ac+ and H3K122ac+ using ChIP-seq data from ENCODE. Expectedly EP300 is enriched across TEs marked with H3K27ac (Fig. 3A). YY1 is enriched at L1s marked with all three acetylations, supporting the known role of YY1 in the activation of the L1 transcription (Athanikar et al., 2004). CTCF and RAD21 showed higher enrichment at H4K16ac+ and H3K122ac+ L1s and LTRs in comparison to H3K27ac+. MYC and KDM1A are depleted at H4K16ac+ and H3K27ac+ L1s. These observations are consistent with the previous reports showing the role of CTCF and RAD21 in activating L1 transcription (Macfarlan et al., 2011; Xu et al., 2014), while MYC and KDM1A repress L1 transcription (Sun et al., 2018). SP1, TCF12 and NANOG binding also specifically enriched at H3K27ac+ L1 and LTRs, suggesting their role in transcription at these elements.

**Fig 3.**
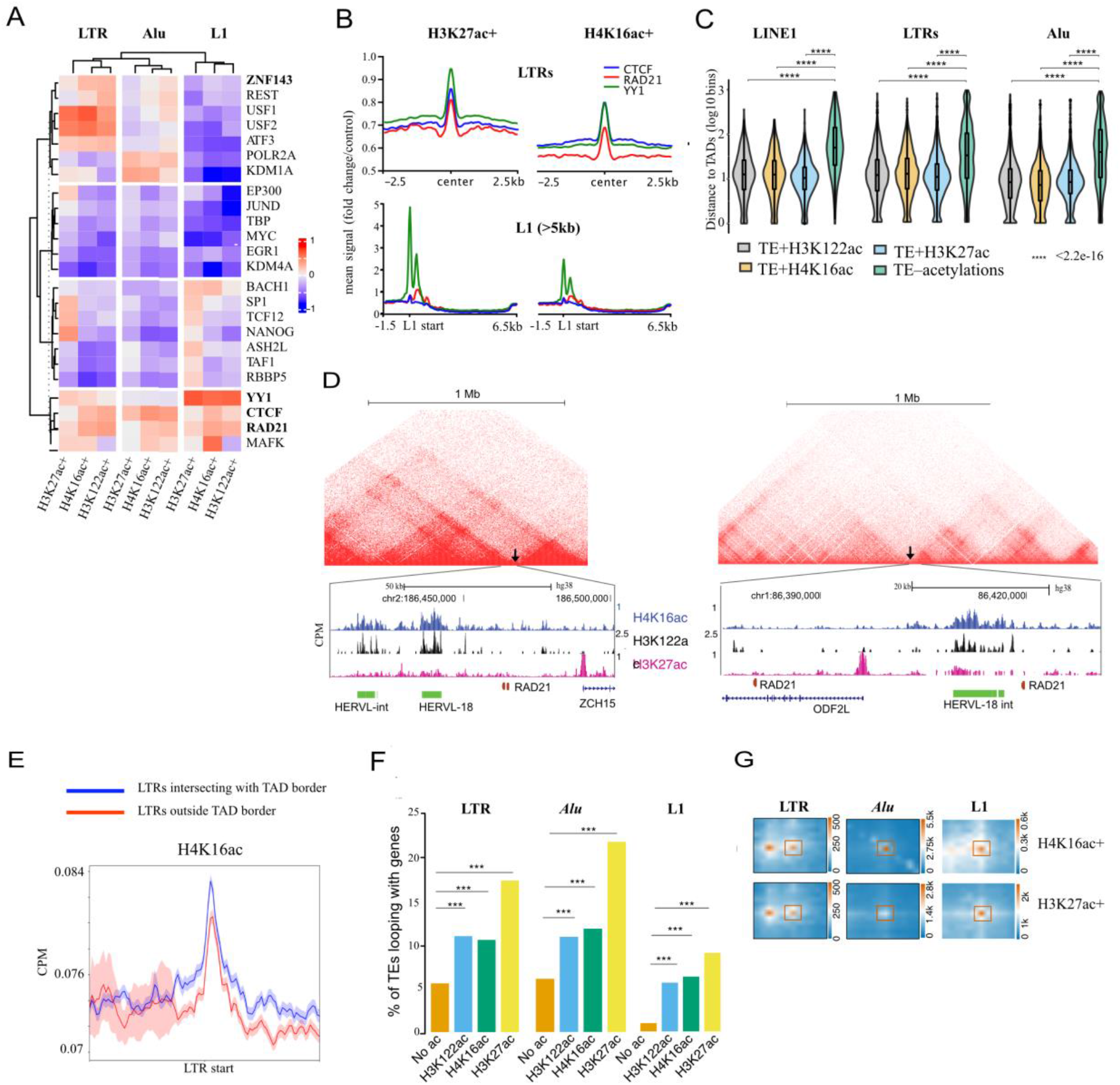
H4K16ac+ L1 and LTRs are enriched at TAD borders and loops with genes. **A**. Heatmap shows the ratio of (difference/sum) for observed and expected occurrences of TF binding sites in H4K16ac, H3K27ac, and H3K122ac peaks at L1 5’ UTR, ERV/LTR and SINE/*Alu* over the random background, a complete list of TFs in fig. S6. **B**. Average-type summary plots showing the mean signal distribution (fold change/control) of YY1 (green), RAD21 (red) and CTCF (blue) at LTR (top) and full-length L1 (>5kb, bottom) that overlaps with H3K27ac (left) or H4K16ac (right). **C**. Violin plot showing the distance to TAD borders (Y-axis, log 10 bins) for LTR, *Alu* and L1 with H3K122ac, H4K16ac, H3K122ac and TEs that lack these acetylations. **D**. Example UCSC genome browser tracks showing H4K16ac, H3K122ac and H3K27ac signals at TAD borders (arrow marks) (micro-C data from H9 hESC). These HERV/LTRs were used for validation by CRISPRi (see Fig. 4D). **E**. Average type summary plot depicting IgG normalised H4K16ac signal (CPM) with standard error (shaded area) at the LTRs overlapping the TAD-border (blue) and LTRs elsewhere in the genome (red). **F**. Bar graph showing the percentage of H4K16ac+, H3K27ac+ and H3K122ac+ and TEs that lack acetylations (full-length L1, LTR and *Alu*) that contact genes via chromatin loops (Fisher’s exact test: ^***^ p-value < 0.001). **G**. Aggregate peak analysis (APA) plots for H4K16ac+ and H3K27ac+ LTR, *Alu* and L1s looping with genes.

### H4K16ac+ L1 and LTRs are enriched at TAD borders and loops with genes

YY1, enriched at acetylated L1s, functions as a looping factor to facilitate interaction between enhancers and promoters (Weintraub et al., 2017). LTRs and *Alu* elements marked with histone acetylations are enriched with USF1, REST and a looping factor ZNF143, compared to L1s marked with acetylations (Fig. 3A and fig. S5). Meta-analysis confirmed the enrichment of CTCF, RAD21 and YY1 at both H3K27ac+ and H4K16ac+ L1 5’ UTRs and LTRs (Fig. 3B). Analysis of published Hi-C data revealed that TEs marked with histone acetylations are enriched at topologically associated domain (TAD) borders than TEs that lack acetylations (Fig. 3C). Moreover, H4K16ac levels are relatively higher at TEs overlapping with the TAD borders than at TEs that do not overlap with TAD borders (Figs. 3D and E). Furthermore, to identify whether TEs with histone acetylations loop with genes, we called significant loops from publicly available micro-C data from H1 hESCs (Krietenstein et al., 2020). This revealed a significantly higher fraction of TEs (L1, LTRs and *Alu*’s) marked with histone acetylations form chromatin loops with genes than TEs that lack acetylations (Figs 3F & G). These analyses show that TEs enriched with histone acetylations contribute to 3D chromatin folding and looping interactions with genes.

### Some of the H4K16ac+ LTR/HERVs act as enhancers to regulate genes in cis

We aimed to investigate the role of H4K16ac+ TEs in the regulation of genes in cis by CRISPR interference (CRISPRi) approach by recruiting of KRAB repressor domain (dCAS9-KRAB) to TEs. We performed CRISPRi for individual HERV/LTR or L1 5’ UTRs, by co-transfecting two independent guide-RNAs that recruit dCAS9-KRAB to specific TEs enriched with H4K16ac in hESCs. We then performed RT-qPCR for nearby expressed genes or genes that show looping interaction in the RAD21 HiChIP data (Fig. 4A) (Lyu et al., 2018). CRISPRi for H4K16ac+ LTR7/HERVH element located ∼50kb away from *PEX1* and ∼30kb from *GATAD1* promoter led to downregulation of *PEX1* but not *GATAD1*(Figs. 4B and D). CRISPRi for another H4K16ac+ LTR7/HERVH element located ∼50kb away from the *NUS1* promoter led to the downregulation of the *NUS1* but not the *GOPC* (Figs. 4C and D). However, CRISPRi for H4K16ac enriched LTRs/HERVL-18, and HERVL-18 int that are close to TAD borders (Figs. 3E and 4D) did not show downregulation of nearby genes *ODF2L* and *ZCF5L*, suggesting some but not all H4K16ac+ HERV/LTRs loci function as enhancers. In contrast, others could contribute to LTR/HERV transcription and 3D genome folding (Figs. 3C-E)(Zhang et al., 2019).

**Fig. 4.**
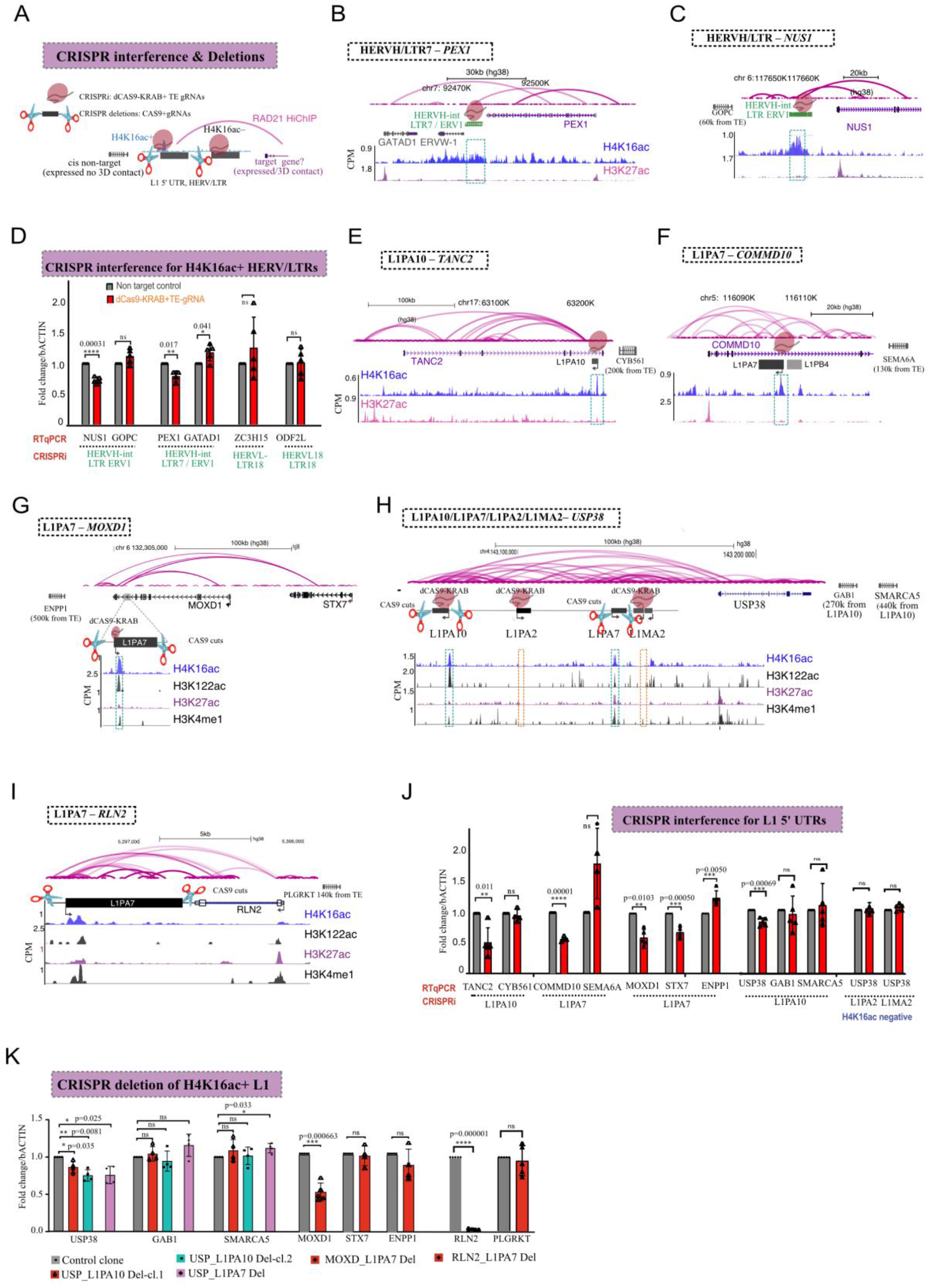
H4K16ac+ L1 5’ UTRs function as enhancers to regulate genes in cis. **A**. Illustration showing CRISPRi and CRISPR-mediated deletion strategy for TEs. Genes that show looping interaction (RAD21-HiChIP data) and are expressed in hESCs are chosen as putative targets for RTqPCR and other nearby expressed genes as controls. **B, C**. Genome-browser tracks showing H4K16ac and H3K27ac CUT&Tag data (CPM) at LTR7/HERVH-int and LTR/ERV1 loci and their putative target genes. **D**. RT-qPCR data showing relative fold change (normalised to b-ACTIN) in the expression of putative target genes *NUS1* and *PEX1* upon CRISPRi for HERV/LTRs but not other nearby genes (GOPC and GATAD1). **E-I**. Like B&C, the genome browser track shows CUT&Tag data at L1PA10 at the *TANC2*, L1PA7 at the *COMMD10*, L1PA7 at the *MOXD1*, L1PA10, L1PA7 at the *USP38* and L1PA7 at the *RLN2* loci. L1PA2 and L1MA7 at the *USP38* locus that lacks histone acetylations were used as controls. **J**. Same as D. but for putative target genes for L1s that are H4K16ac+ and L1s that are H4K16ac– (L1PA2 and L1MA2) TANC2, COMMD10, MOXD1/STX7 and USP38 were selected as putative target genes along with CYB561, SEMA10A, ENPP1, and GAB1/SMARCA5 were selected as putative non-targets. **K**. Same as J, but RTqPCR upon CRISPR CAS9 mediated deletion of full-length L1. Two independent clones for L1PA10 and one for L1PA7 located at the USP38 locus were tested, and for L1PA7 located at MOXD1 and RLN2, the pools of cells were tested.

### H4K16ac+ L1 5’ UTRs function as enhancers to regulate genes in cis

We next focused on L1s and asked whether L1 5’ UTRs enriched with H4K16ac regulate genes in cis by performing CRISPRi for H4K16ac+ L1 5’ UTRs, together with two L1 5’ UTRs that lack detectable histone acetylations. CRISPRi for H4K16ac+ 5’ UTR of an L1PA10 located ∼110kb upstream of *USP38* led to specific downregulation of *USP38* but not other nearby genes *GAB1* and *SMARCA5* (Figs. 4E and J). Notably, CRISPRi for two 5’ UTRs of L1’s that lack H4K16ac located ∼30kb and ∼85kb from the *USP38* promoter led to no change in USP38 transcripts level, showing the specificity of H4K16ac+ L1PA10 element in regulating *USP38*. Similarly, CRISPRi for H4K16ac+ 5’ UTR of L1PA10 located ∼270kb from the *TANC2* promoter, led to downregulation of *TANC2*, but not *CYB561* (Figs. 4F and J). CRISPRi for H4K16ac+ L1PA7 located ∼24kb from *COMMD10* promoter also led to a significant downregulation of *COMMD10*, but not the nearby gene *SEM6A* (Figs. 4G and J). RAD21 HiChIP data and the micro-C analysis revealed the significant looping interactions between *MOXD1* and *STX7* genes with the H4K16ac+ L1PA8, located ∼100kb away from the *MOXD1* promoter (Fig. 4H) (data S1). CRISPRi for 5’ UTR of this L1PA8 led to significant downregulation of both *MOXD1* and *STX7*. However, the expression of *ENPP1*, which does not loop with this L1, is not altered, demonstrating the specific cis-regulatory function of these L1s (Fig. 4J).

To further confirm that H4K16ac+ L1 5’ UTRs regulate genes in cis, we utilised CRISPR CAS9 to delete four full-length L1 elements in h1 hESCs (∼7 kb). Due to the difficulty in specific deletion of L1 5’ UTRs, we nucleofected CAS9 with pairs of synthetic guide RNAs that target the flanking region of four full-length L1s. We generated two independent clonal lines with heterozygous deletions for L1PA10 and one clone for L1PA7; both are H4K16ac+ and located upstream of *USP38* (Fig. 4H and fig. S9). Consistent with CRISPRi data, RT-qPCR data showed that deletion of L1PA10 and L1PA7 led to the downregulation of *USP38* but not other tested nearby genes *GAB1* and *SMARCA5* (Fig. 4K). For deletion of L1s located at the *MOXD1* and *RLN2* loci (Fig 4G and I), we nucleofected gRNA/CAS9 ribonucleoprotein complexes and used two independent pools of hESCs that showed ∼50% deletion efficiency (fig. S6) for RT-qPCR. Although CRISPRi for L1PA8 resulted in the downregulation of both *MOXD1* and *STX7*, genetic deletion led to specific downregulation of *MOXD1* but not *STX7* and *ENPP1* (Fig. 4H and 4K). Deletion of another H4K16ac+ L1PA7 located downstream of *RLN2*, ∼12 kb away from the *RLN2* promoter, led to the downregulation of *RLN2* but not a nearby gene *PLGRKT* (Fig. 4I and 4K). Overall, CRISPR interference and genetic deletions experiments confirmed that H4K16ac+ L1 and LTRs contribute to the transcriptional activation of genes in cis.

### MSL-mediated H4K16ac regulates the transcription of L1 and ERV/LTRs

Next, we aimed to investigate whether H4K16ac regulate TE transcription by depleting H4K16ac. H4K16ac is catalysed explicitly by KAT8 when associated with male-specific lethal (MSL), but not non-specific lethal (NSL) complex (Chatterjee et al., 2016; Chelmicki et al., 2014; Radzisheuskaya et al., 2021; Ravens et al., 2014) (Fig. 5A). Since depletion of individual MSL complex proteins MSL1, MSL2 and MSL3 are sufficient to reduce H4K16ac level (Monserrat et al., 2021). We depleted MSL3 using two independent lentiviral shRNAs in H9 hESCs; we first validated the depletion by RTqPCR, which showed ∼50% downregulation of MSL3. RT-PCR with primers recognising full-length L1 subfamilies such as human-specific (L1HS), mammalian-wide (L1M) and primate-specific (L1PA and L1PB) full-length L1s showed significant downregulation upon MSL3 knockdown (KD). Similarly, RT-qPCR with primers recognising HERV-K and HERV-H transcripts showed significant downregulation in MSL3-depleted hESCs (Fig. 5B). Western blotting confirmed MSL3 depletion led to a specific reduction in H4K16ac but not H3K27ac (Fig. 5C). Like the transcripts data, L1-ORF1 and HERV envelope protein (antibody raised against ERVW-1) were also reduced upon MSL3 and H4K16ac depletion (Fig. 5C), consistent with the high level of H4K16ac at ERVW-1 locus (Fig. 4B).

**Fig. 5.**
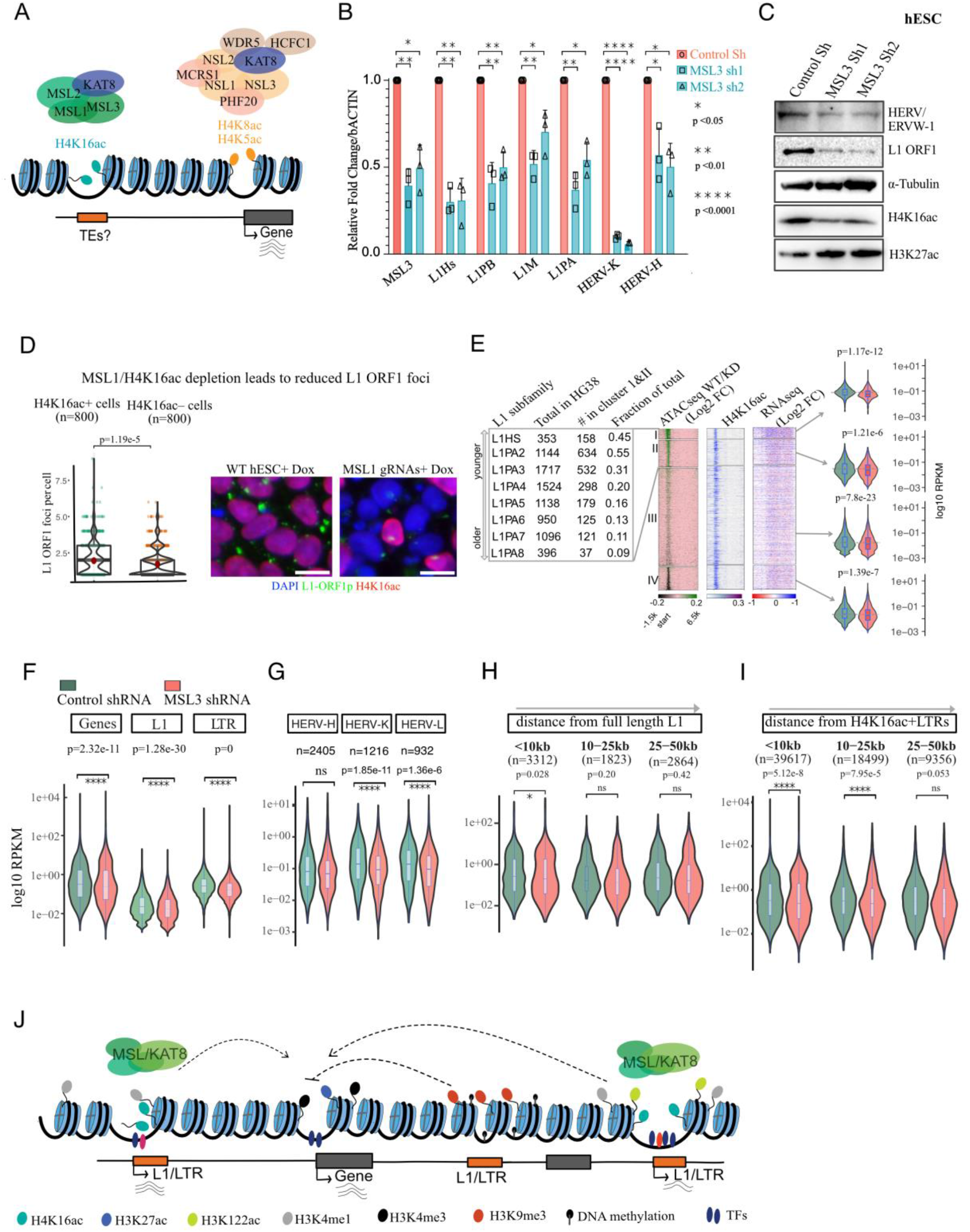
MSL regulate the transcription of TEs and genes in cis. **A**. Illustration showing KAT8 catalyses H4K16ac only when it is bound to MSL and but not NSL complex. **B**. RT-qPCR data from hESCs showing mean ± SD (n=3) fold change (normalised to β-ACTIN) in MSL3, L1 and HERV subfamilies upon lentiviral shRNA mediated knockdown of MSL3 using two independent shRNAs versus non-target control. **C**. Like B, Western blots showing HERV, L1-ORF1 and H4K16ac levels, α-Tubulin and H3K27ac served as controls in control and MSL3 knockdown hESCs. **D**. Representative images (right) and quantification of high-content imaging data showing the number of L1 ORF1p foci per cell (n=800) (left) in H4K16ac+ and H4K16ac– MSL1 knockout cells. **E**. Heatmaps showing H4K16ac, ATAC-seq and RNAseq (log2 fold change) for control/MSL3 knockdown in hESCs, n=4) across full-length L1 with K-means clusters. The distribution of L1 subfamily members in clusters 1 and 2 (left). Violin plots (right) showing RNAseq signal (log10 RPKM, control and MSL3 knockdown in hESCs) at four L1 clusters. Boxes indicate the median and interquartile range, with whiskers showing the first and the fourth quartile. **F**. Violin plots showing RNAseq for genes, full-length L1s and LTRs for control and MSL3 knockdown in hESCs (n=4). **G**. Like F, RNAseq signal at HERVs subclasses HERV-K, HERV-H and HERV-L. **H**. Like F and G, RNAseq signal at genes that lie in the 10kb, 10-25kb or 25-50kb distance from the H4K16ac overlapping full-length L1s. **I**. Like H, RNAseq signals at genes that lie in the 10kb, 10-25kb or 25-50kb distance from the H4K16ac overlapping LTRs. Statistical analyses for all the violin plots were done using the Dunn test with Bonferroni correction. **J**. The working model shows MSL/KAT8 mediated H4K16ac maintains accessible chromatin, activates transcription at TEs and contributes to their enhancer activity to regulate genes in cis.

We further used doxycycline-inducible Cas9 (iCAS9) mediated knockout (KO) of MSL1 in H1 hESCs (Fig. 5D) and MSL1 and MSL3 KO in TDFs (fig. S7) to confirm our findings from the shRNA mediated MSL3 depletion. Immunofluorescence for H4K16ac and L1-ORF1 protein followed by high-content imaging revealed a significantly reduced number of L1-ORF1p foci in H4K16ac depleted MSL1 KO hESCs (Fig. 5D). Like hESC data, MSL3 KO in TDFs reduced the bulk of H4K16ac (fig. S7A) and at L1 5’ UTRs and LTRs (fig. S7D).

MSL3 KD RNA-seq analysis in hESCs showed that pluripotency and differentiation-associated genes were unaffected (fig. S8A and B). However, H4K16ac+ genes were more affected than H4K16ac-genes (fig. S8C). Further analysis on L1s and LTRs showed significant downregulation of both human-specific (L1HS), primate-specific (L1PA2 to L1PA16) full-length L1 and LTR sub-family transcripts (Fig. 5E and fig. 5B). L1 and HERV-K and HERV-L transcripts and protein-coding genes also show small but significant downregulation in MSL3 KD (Figs. 5F. G and fig. S8D). However, RNAseq data analysis from MSL3 KO TDFs showed significant downregulation of L1 and LTR transcripts (fig. S7B and C). Notably, H4K16ac+ L1s but not H4K16ac– L1s are significantly downregulated in MSL1 KO TDFs (p=0.000104) (fig. S7B). All these results confirm the direct role of MSL/H4K16ac in the transcriptional activation of L1.

### MSL/H4K16ac at L1/LTRs maintain active cis-regulatory landscape

H4K16ac causes chromatin decompaction *in vitro*, and depletion of H4K16ac has been shown to reduce chromatin accessibility (Samata et al., 2020; Shogren-Knaak et al., 2006). Therefore, we asked whether the lack of H4K16ac leads to altered accessibility at TEs. ATAC-seq data showed a specific reduction in accessible DNA at the 5’ UTR of L1s in MSL3-depleted hESCs (Fig. 5E). Many evolutionarily younger L1s show a strong reduction in DNA accessibility accompanied by reduced transcriptional activity at these elements (Fig. 5E).

We next asked whether depletion of MSL/H4K16ac at L1 and LTRs affects the expression of genes associated with H4K16ac+ TEs. MSL3 depletion led to a small but significant downregulation of many transcripts (n=3,312) closer (<10 kb) to H4K16ac+ L1s (Fig. 5H). Similarly, many transcripts that are closer (<10 and <25 kb) to H4K16ac+ LTRs are significantly downregulated compared to transcripts that are away (Fig. 5I). Overall, our results confirm the role of MSL/H4K16ac at L1 and LTRs not only in transcriptional activation of TEs (Fig. 5B, D) but also in regulating genes that they are associated in linear distance or 3D space (Figs 4 and 5H, I). Therefore, we conclude that MSL/KAT8-mediated H4K16ac leads to the opening of chromatin structure and increased transcriptional activity at L1 and LTRs in a cell type-specific manner. The permissive local chromatin environment at H4K16ac marked TEs shapes the cis-regulatory landscape across the mammalian genome.

## Discussion

TEs are repressed by many epigenetic pathways, such as DNA methylation, H3K9me3 and KRAB-ZNF and HUSH complex, and piRNAs. We have discovered that MSL/H4K16ac axis functions as a transcriptional activator of TEs, particularly L1s and LTRs. TEs have contributed significantly to the evolution of mammalian genomes by helping to shape both the coding and non-coding regulatory landscape. Some LTR subfamilies are known to function as enhancers. Here for the first time, we have demonstrated that L1 5’ UTRs and LTR/ERVs enriched with histone acetylations loop with genes and L1 and LTRs marked with H4K16ac function as enhancers to regulate genes in cis.

Roadmap epigenomics data showed that TEs are depleted of H3K27ac and accessible chromatin and only 3% of TE bases are annotated with active regulatory chromHMM states, compared to 32% of promoter bases (Pehrsson et al., 2019). However, TEs contribute up to 40% of TF binding sites; hence TEs are proposed to contribute to species and tissue-specific rewiring of gene regulatory networks (Kunarso et al., 2010; Sundaram et al., 2014, 2017). This suggest the unknown chromatin pathway that could contribute to enhancer activity of TEs in a cell type or species-specific manner, which could be independent of H3K27ac. Our CUT&Tag data shows the level of H3K27ac is much higher at genes and promoters compared to TE. However, H4K16ac is enriched explicitly at the L1 5’ UTRs, along with several other chromatin features associated with active enhancers. We now demonstrate that L1 5’ UTRs marked with H4K16ac along or together with H3K122ac and H3K27ac function as enhancers to regulate the expression of genes in cis. Although L1s are expressed at higher levels during early development, including stem cells, they are also upregulated in the neuronal lineage. Consistently, we found that H4K16ac is enriched at L1 5’ UTRs in human and mouse stem cells, cancer cell lines and post-mortem brain tissues. Suggesting that 5’ UTRs of L1s bound by tissue-specific TFs and enriched with histone acetylations could function as tissue-specific enhancers. Enrichment of H4K16ac at TEs, which constitute a significant part of the mammalian genome, is consistent with the findings showing that nearly 30% of the histone H4 is acetylated at H4K16(Radzisheuskaya et al., 2021).

Many LTR subfamilies are enriched with chromatin features that predict them as active enhancers and are suggested to be essential to drive the expression of lineage-specific genes(Fuentes et al., 2018; Kunarso et al., 2010). However, only a minority of putative RLTR13D6 subfamily-derived enhancers identified through epigenomic analyses were experimentally validated to function as enhancers (Todd et al., 2019). This highlights the importance of functional validations using CRISPR-based perturbation of candidate TEs enriched with enhancer chromatin features. Although we found all of the tested CRISPR-edited H4K16ac+ L1s downregulated their target genes in cis. Genome-wide enhancer reporter assays, in combination with systematic genome-scale perturbation, are needed to identify what fraction of L1 and LTRs with H4K16ac function as enhancers.

TEs marked with acetylations, including H4K16ac, are bound by looping factors, including CTCF, RAD21, YY1 and ZNF143. Moreover, a significantly higher fraction of these TEs loops with genes compared to TEs that lack acetylations, further supporting the role of transcriptionally active TEs in rewiring the regulatory landscape in species and cell type-specific manner. This is consistent with the previous report showing transcriptionally active L1 and HERVs contribute to TAD boundary formation (Zhang et al., 2019). Since our results show MSL/H4K16ac axis drives the transcription at TEs, including HERVs (Fig. 5 and E, fig. S4B), we hypothesise that MSL/H4K16ac-mediated transcription at TEs likely contributes to the rewiring of 3D chromatin organisation at transcriptionally active TEs. The factors contributing to the recruitment of MSL complex to the specific genomic region are unknown in mammals. Intriguingly, however, the role of MSL complex to co-opt TEs to rewire cis-regulatory elements appears to be conserved during the evolution of dosage compensation in *Drosophila miranda*, where a mutant Helitron TE is shown to recruit MSL complex to evolutionarily young X chromosome to increase transcription (Christopher and Bachtrog, 2013). TFs enriched at H4K16ac+ TEs (Fig. 3A, fig. S5A) could contribute to maintaining MSL/H4K16ac and transcription at TEs. Notably, the MSL complex recruits YY1 to the *Tsix* promoter to activate its expression in mESCs (Chelmicki et al., 2014), suggesting a possible interplay between MSL and YY1 in regulating L1 transcription. Interestingly, MAFK, which was previously reported to be enriched at TEs (Sundaram et al., 2014), is enriched explicitly at H4K16ac+ L1 5’ UTRs, suggesting a potential interplay between MAFK and MSL complex.

Neuronal cells have high L1 expression and retrotransposition (Macia et al., 2017); retrotransposon dysregulation is also linked with neurological disorders (Hancks and Kazazian, 2016). Loss of function mutations in genes encoding KAT8 containing protein complexes such as *KANSL1, MSL3*, and *KAT8* lead to neurodevelopmental disorders (Basilicata et al., 2018; Koolen et al., 2012; Sharp et al., 2006; Shaw-Smith et al., 2006). Enrichment of H4K16ac at the 5’ UTRs of L1s in human brain tissues suggests that the TE dysregulation in the nervous system could be a possible mechanism for these disorders (fig. S3) (Nativio et al., 2018). Further studies on the specific role of H4K16ac in neuronal cell types will reveal whether H4K16ac dysregulation could contribute to neuronal-specific dysregulation of TEs and gene expression, contributing to neurodevelopmental and neurodegenerative disorders.

In yeast, H4K16ac regulate life span and cellular senescence (Dang et al., 2009). Senescent cells show enrichment of H4K16ac to promoter regions of expressed genes (Rai et al., 2014). Analysis of publicly available H4K16ac ChIP-seq data showed a dramatic loss of H4K16ac across L1s and LTRs in senescent compared to proliferating cells (fig. S9), suggesting that proliferating cells have adapted to permissive chromatin at TEs compared to replicative senescent cells. However, in contrast, L1 elements become transcriptionally derepressed during cellular senescence and activate the interferon I (IFN-I) response (De Cecco et al., 2019). Further investigation will be needed to understand the direct role of the H4K16ac pathway in regulating L1 transcription during senescence.

In summary, we show that H4K16ac marked L1 and LTRs act as enhancers to regulate genes in cis. The act of transcription at L1 5’ UTRs and LTRs mediated by H4K16ac could contribute to chromatin topology and enhancer-mediated regulation of host gene expression in cis, as L1 and LTRs that are marked with histone acetylations are located within the regulatory elements, or they interact with genes. The permissive chromatin structure mediated by H4K16ac and H3K122ac could counteract the epigenetic repressive environment at the TEs within the regulatory elements (Fig. 5J) (Liu et al., 2018).

## Acknowledgements

We thank QMUL epigenetics hub members Alex de Mendoza, Chris Bell, Vardhman Rakyan and Lovorka Stojic for discussing and reading the manuscript. We thank Ludovic Vallier (Cambridge UK, with MTA from WiCell) for sharing the H9 cell line and Steve Henikoff (Fred Hutchinson Cancer Research Center) lab for pATn5 expression plasmid. We thank the labs of Edda Schulz (Max Planck Institute for Molecular Genetics) and Chema Martin (Queen Mary University of London) for sharing purified pATn5. Pankaj Dubey, Ivan Alic and Aoife Murrey help in hESC cell culture. We thank Garry Warnes and Luke Gammon from Blizard Institute core facilities for help in flow sorter and high content imaging analysis. This research used Apocrita HPC, supported by QMUL Research-IT. For Open Access, the author has applied a CC BY public copyright license to any Author Accepted Manuscript version arising from this submission.

## Author contributions

Conceptualisation: MMP, DP, MP

Methodology: MMP, DP, MP, FB, JS, MRB, PS, OAG, NRZ, SS

Investigation: MMP, DP, MP, FB, JS, SB

Visualisation: MMP, DP, MP, FB, MRB

Funding acquisition: MMP

Project administration: MMP Supervision: MMP

Writing – original draft: MMP, DP, MP

Writing – review & editing: MMP, DP, MP, FB, JS, SB, NRZ, MRB

## Data and materials availability

The data discussed in this publication have been deposited in NCBI’s Gene Expression Omnibus (Pal et al., 2022) and are accessible through GEO Series accession number GSE200770 (https://www.ncbi.nlm.nih.gov/geo/query/acc.cgi?acc=GSE200770). All the datasets generated and used in this study are detailed in Table S1.

