## Supplementary material for "H4K16ac activates the transcription of transposable elements and contributes to their cis-regulatory function": H4K16acSuppl.file15112022Biorxiv.pdf

### This PDF file includes:

Online Methods
Supplementary Figs. 1 to 9
Supplementary Table 1 to6
Supplementary references

### Online Methods

#### Cell Culture and Transduction

The H9 hESC line was a kind gift from Ludovic Vallier's lab with the MTA from WiCell. H1 hESCs expressing doxycycline-inducible CAS9 (H1 iCAS9) cells were generated in Silvia Santos lab (The Crick Institute). hESCs were grown on geltrex coated plates (ThermoFisher Scientific, A1413302) in mTeSR™ Plus medium (StemCell Technologies, #100-0276) supplemented with 100 U/ml penicillin-streptomycin (Gibco, 15140122) and passaged every 3-4 days with ReLeSR™ (StemCell Technologies, 100-0484) according to manufacturer's protocols.

Transformed dermal fibroblasts (TDF) expressing guide RNAs (3 guides per pool) targeting MSL1 and MSL3 and parental (WT) TDF lines were generated in Paola Scaffidi's lab (The Crick Institute). Cells were grown in MEM (Gibco, 11095080) supplemented with 10% FBS (Sigma, F7524), 1X Glutamax (Gibco, 35050061), 1X non-essential amino acid solution (Sigma, M7145) and 100 U/ml penicillin-streptomycin.

iCAS9 cells were transduced with three lentiviral guide RNAs targeting MSL1 and MSL3(Monserrat et al., 2021). Parental iCAS9 H1, iCAS9 with MSL guide RNAs and TDF iCas9 transduced with MSL1, and MSL3 guide RNA pools were treated with 1μg/ml doxycycline (Sigma) to generate the inducible MSL KO lines[27]. After 4 to 7 days of doxycycline induction, the knockout was validated by immunofluorescence followed by high-content microscopy and western blot using H4K16ac and H3K27ac antibodies.

HEK293T cells were grown in DMEM, high glucose (Lonza, BE12-614Q) supplemented with 10% FBS (Sigma, F7524), 1X Glutamax (Gibco, 35050061) and 100 U/ml penicillin-streptomycin. HEK293 and HeLa cells were grown in DMEM, high glucose (Lonza, BE12-614Q) supplemented with 10% FBS (Sigma, F7524), 1X Glutamax (Gibco, 35050061) and 100 U/ml penicillin-streptomycin. PC3 and LNCaP cells were grown in RPMI medium (Gibco, 21875034) supplemented with 10% FBS (Sigma, F7524) and 100 U/ml penicillin-streptomycin. RWPE1 cells were grown in a keratinocyte serum-free medium (Gibco, 10724011) supplemented with 100 U/ml penicillin-streptomycin. K562 cells were grown in Iscove's Modified Dulbecco's Medium (Lonza, BE12-722F) supplemented with 10% FBS (Sigma, F7524) and 100 U/ml penicillin-streptomycin. SH-SY5Y cells were grown in DMEM / F12 (1:1) medium (Gibco, 11320033) supplemented with 10% FBS (Sigma, F7524) and 100 U/ml penicillin-streptomycin. All the cell lines were tested for Mycoplasma contamination using EZ-PCR Kit (Geneflow, K1-0210).

For the generation of MSL3 stable knockdown H9 hESCs, cells were transduced with lentiviral particles (Sigma, Mission shRNAs, MSL3 sh1 TRCN0000022105, MSL3 sh2 TRCN0000022107 and mammalian non-targeting shRNA (SHC002V) at an MOI of 6. 48 hours after transduction, cells were selected with 0.5 μg/ml puromycin (Gibco, A1113803) for 48 hours and surviving cells were then allowed to recover until they formed viable colonies.

#### Western blotting

Cells were pelleted by centrifugation at 228 *g* for 5 min at 4°C and resuspended in RIPA buffer (150 mM sodium chloride, 1.0% NP-40 or Triton X-100, 0.5% sodium deoxycholate, 0.1% SDS (sodium dodecyl sulfate) and 50 mM Tris, pH 8.0) and protease inhibitors + Benzonase (Novagen; final concentration, 1.25 U/μl) and incubated for 30 min on ice with intermittent mixing. Extracts were sonicated for 5 cycles with Bioruptor (Diagenode) with the 30s on and 30s off cycles and cleared by centrifugation at 15500 *g* for 10 min at 4°C. Equal amounts of protein extract were denatured in 1x Bolt LDS sample buffer (ThermoFisher Scientific, B0007) and separated on Bolt Bis-Tris gels (ThermoFisher Scientific, NW04120BOX, NW00122BOX), blotted to Polyvinylidene fluoride (PVDF) membrane (BioRad, 1704156) and immunoblotted with antibodies recognising; MSL3 (Merck Millipore, ABE467, 1:1000 dilution), L1 ORF1 (Merck Millipore, MABC1152, 1:1000 dilution), H4K16ac (Abcam, ab109463, 1:5000 dilution), H3K27ac (Abcam, ab4729, 1:5000 dilution), α-tubulin (Sigma, T9026, 1:5000 dilution) and HERV (Novus Biologicals, NB100-93579, 1:500 dilution), goat anti-rabbit IgG H&L HRP (Abcam, ab6721) and goat anti-mouse H&L HRP (ThermoFisher Scientific, 31430) secondary antibodies.

**Immunofluorescence and imaging**

Cells were grown on 24 well cell culture plates, fixed with 4% formaldehyde, and incubated for five minutes with permeabilisation buffer (PBS containing 0.1% triton X100), block with PBS containing 0.1% Triton X 100 and 2% BSA) for one hour. Primary antibodies, H4K16ac (Abcam, Ab109463, 1:500) and L1 ORF1 (Merck Millipore, MABC1152, 1:500 dilution) overnight at 4C, washed three times with PBS (10 minutes each) and incubated with anti-rabbit secondary antibodies (Abcam, Ab150080, 1:500) and DAPI (1:1000). After washing three times with PBS (10 minutes each), the cells were left in PBS and imaged with Incell2000.

**CUT&Tag**

CUT&Tag was performed according to Kaya-Okur et al (Kaya-okur et al., 2019) protocol with modifications to tissue processing as described below. Experiments were performed in biological duplicates from each cell type. ~100,000 cells were pelleted by centrifugation for 3 min at 600 x g at room temperature and resuspended in 500μl of ice-cold NE1 buffer (20 mM HEPES-KOH pH 7.9, 10 mM KCl, 0.5 mM spermidine, 1% Triton X-100, and 20 % glycerol and cOmplete EDTA free protease inhibitor tablet) and was let to sit for 10 min on ice. Nuclei were pelleted by centrifugation for 4 min at 1300 x g at 4°C, resuspended in 500μl of wash buffer and held on ice until beads were ready. The required amount of BioMag Plus Concanavalin-A-conjugated magnetic beads (ConA beads, Polysciences, Inc) were transferred into the binding buffer (20 mM HEPES-KOH pH 7.9, 10 mM KCl, 1mM $CaCl_2$ and 1mM $MnCl_2$), washed once in the same buffer, each time placing them on a magnetic rack to allow the beads to separate from the buffer and resuspended in binding buffer. 10 μl of beads was then added to each tube containing cells and rotated on an end-to-end rotator for 10 minutes. After a quick spin to remove liquid from the cap, tubes were placed on a magnet stand to clear, the liquid was withdrawn, and 800μl of antibody buffer containing 1μg of primary antibodies: normal rabbit IgG (Santa Cruz Cat no sc-2027), H3K27ac (Abcam, ab4729), H4K16ac (Abcam, ab109463), H3K122ac (Abcam, ab33309), H3K4me1(Abcam, ab8895), H3K36me3 (Abcam, ab9050)) H3K4me3 (Millipore, 07-473), H3K27me3 (Abcam, ab192985) and H3K9me3 (Abcam, ab176916)) was added and incubated at 4ºC overnight in a nutator. Secondary antibodies (guinea pig α-rabbit antibody, Antibodies online, ABIN101961) were added 1:100 in Dig-wash buffer (5% digitonin in wash buffer), and 100 µL was squirted in per sample while gently vortexing to allow the solution to dislodge the beads from the sides and incubated for 60 min on a nutator. Unbound antibodies were washed in 1 ml of Dig-wash buffer three times. 100 µl of (1:250 diluted) protein-A-Tn5 loaded with adapters in Dig-300 buffer (20 mM HEPES pH 7.5, 300 mM NaCl, 0.5 mM spermidine with Roche cOmplete EDTA free protease inhibitor) was added to the samples, placed on nutator for 1 hour and washed three times in 1 ml of Dig-300 buffer to remove unbound pA-Tn5. 300 µL Tagmentation buffer (Dig-

300 buffer + 5 mM $MgCl_2$) was added while gentle vortexing, and samples were incubated at 37ºC for 1 hr on an incubator. Tagmentation was stopped by adding 10 µL 0.5M EDTA, 3 µL 10% SDS and 2.5 µL 20 mg/mL Proteinase K to each sample. Samples were mixed by full-speed vortexing for ~2 seconds and incubated for 1 hr at 55ºC to digest proteins. DNA was purified by phenol: chloroform extraction using phase-lock tubes (Quanta Bio) followed by ethanol precipitation. Libraries were prepared using NEBNext HiFi 2x PCR Master mix (M0541S) with a 72ºC gap-filling step followed by 13 cycles of PCR with 10-second combined annealing and extension for the enrichment of short DNA fragments. Libraries were sequenced in Novaseq 6000 (Novogene) with 150bp paired-end reads.

**RT-qPCR**

Total RNA was isolated from H9 hESCs using TRIzol reagent (ThermoFisher Scientific, 15596026). For reverse transcriptase-polymerase chain reaction (RT-qPCR), cDNAs were prepared with LunaScript® RT SuperMix Kit (NEB, E3010). For CRISPRi experiments, RNA isolation was done using a kit (Monarch, T2040S) followed by reverse transcription using LunaScript® RT SuperMix Kit (NEB, E3010), qPCR using qPCRBIO SyGreen Mix Lo-ROX (PCRBio) in LightCycler 480 instrument (Roche). The list of specific primers used is given in Supplementary Table 3. RTqPCR was done with three independent biological replicates, each of control shRNA and two independent shRNAs targeting MSL3 or relevant empty vector controls and dCAS9 systems for CRISPRi, on a StepOnePlus™ Real-Time PCR System (Applied Biosystems). Data were normalised to β-actin from three biological replicates.

**RNA sequencing**

RNA was isolated using Monarch RNA mini prep kit (NEB) with genomic DNA elimination column and on-column DNase treatment. MSL3 KD RNA sequencing libraries were prepared by spiking in equal amounts of The External RNA Controls Consortium (ERCC) Spike-in RNA Variant Control Sets (SIRV set 3, Lexogen), and 500 ng of RNA was used for depletion of rRNA using RiboCOP kit (Lexogen), followed by RNAseq library preparation using CORALL Total RNA-Seq Library Prep Kit (Lexogen). Libraries were sequenced as 150 bp paired-end reads using Novaseq 6000. In the case of H1 iCAS9 and MSL1 KO RNAseq, Ribosomal RNAs were depleted using NEBNext® rRNA Depletion Kit (Human/Mouse/Rat) (NEB #E7400) followed by library preparation using NEBNext® Ultra™ II Directional RNA Library Prep Kit for Illumina® (NEB #E7765).

**ATAC-seq**

ATAC-seq is performed as described in (Buenrostro et al., 2013) with the modifications. The freshly harvested 50,000 cells were washed in PBS and resuspended in a resuspension buffer (10mM Tris-HCl, 10mM NaCl, 3mM MgCl2). Cells were resuspended and incubated on ice for 3 min in 50ul of cold lysis buffer (0.1% NP-40, 0.1% Tween-20, 0.01% Digitonin in resuspension buffer). Nuclei were washed in 1 ml of wash buffer (990ul resuspension buffer, 0.1% Tween-20) by inverting three times. Nuclei were pelleted by centrifuging at 500x g for 10 min at 4C. The nuclei were resuspended in 47.5 ml of Nextera Tagmentation buffer (Nextera DNA Sample Preparation Kit) and incubated with 2.5 µl of the Tn5 transposase (Nextera kit, Illumina) at 37°C for 30 min. The resulting DNA fragments were purified using a miniElute column (Qiagen) and amplified by NEBNext High-Fidelity PCR Master Mix in a total volume of 50 µl. The thermocycling protocol for this reaction was 72°C for 5 min, 98°C for 30 s and five cycles of 98°C for 10 s, 63°C for 30 s and 72°C for 1 min. Using the universal adapter primer and a unique barcoded adapter primer (same as CUT&Tag primers). To avoid over-amplification, after the initial five cycles, the number of remaining cycles required was estimated for each sample using qPCR by adding SYBRGreen and using 5 µl of the previous PCR as a template. The number of additional cycles was determined to be the number it took for the qPCR to reach one-third of maximal fluorescence. The original PCR was then resumed, and each sample was cycled as necessary. After amplification, the samples were purified using AMPure XP beads. The libraries were sequenced as a Minimum of 50 million 150 bp paired-end reads in Novoseq (Novogene PLC).

**CRISPRi with dCAS9-KRAB**

CRISPRi using dCAS9-KRAB is performed as described in (Pradeepa et al., 2016) with the following modifications. The CRISPR-Bac plasmid (PB_tre_dCas9_KRAB, Addgene ID 126030) (Schertzer et al., 2019), a kind gift from J. Mauro Calabrese, was mixed with the piggyBac-transposase plasmid in a 1:1 ratio (2 µg each for one well of a 6-well) into opti-MEM along with TransIT-LT1 in a 1:3 ratio (Mirus, MIR2300) and reverse transfected into H9 hESCs according to manufacturer's protocol. The next day, the cells were allowed to recover from the transfection for 24 hours and then selected with 100 µg/ml hygromycin B for 5 days. Surviving colonies were then expanded and reverse-transfected with various guide RNA expressing plasmids (cloned into pSLQ1371) with TransIT-LT1. 1.25X10^6 cells were reverse transfected with 1 µg of the guide RNA expressing plasmid (per well of a 24-well plate). To improve the efficiency of plasmid delivery, the transfection was repeated the next day (forward transfection). 48 hours after the first transfection, cells were briefly selected with puromycin (0.5 µg/ml) for 24 hours and left to recover for 96 hours from the first transfection, cells were harvested for RNA isolation and RT-qPCR.

**CRISPR–Cas9 deletion of LINE1 elements in hESCs**

Two crRNAs performed LINE1 element deletions were designed to target nonrepetitive flanking sites of the LINE1 elements (Table S5). Individual crRNAs were mixed with tracerRNAs Alt-R® CRISPR-Cas9 tracrRNA, ATTO 550, to and mixed with CAS9 protein (Alt-R® S.p. HiFi Cas9 Nuclease V3) to form ribo-nucleocomplex, 200K per well H1 hESCs were nucleofected in the presence of Alt-R® Cas9 Electroporation Enhancer in 16 strips format using primary cell kit (P3). hESCs were electroporated using a 4D nucleofector, the P3 Primary Cells 4D-Nucleofector X kit S (Lonza, LOV4XP3032), with the pulse program. After nucleofection, cells were resuspended in an hESC medium supplemented with ROCK inhibitors and seeded to geltrex-coated 96 wells for 2 days at 37 °C in a humidified incubator with 5% $CO_2$. hESCs were split into 96 wells and 6 well plates to pick single-cell colonies. The pool of cells 5 days after nucleofection were harvested for checking the deletion efficiency and RT-qPCR. Cells were seeded in 6 well plates for picking single cell colonies; the deletion was assessed by rapid DNA lysis and PCR using PCRBIO Rapid Extract PCR kit (PB10.24-40). PCR products were subjected to Sanger sequencing. Primer sequences used for screening are listed in Table S4.

**Analysis of CUT&Tag-seq data**

**Mapping**

150bp paired-end reads for the CUT&Tag-seq were trimmed for adapters using the Trimmomatic tool and aligned to the hg38 genome through local Bowtie2 (version 2.4.5) with these parameters for pair-end mapping: *--very-sensitive-local --no-unal --no-mixed --no-discordant --phred33 -I 10 -X 700*. For analyses, multi-mapped reads were filtered out and only uniquely mapped reads were retained with the samtools flag of *-q 2 -f 0x200*. For individual replicates, the bam files were sorted, indexed and used for generating bedgraphs (for peak calling) and bigwigs. The bam files were sorted and indexed using the *samtools* (version 1.9) *sort* and the samtools *index*. Merging of multiple replicates was performed using *samtools merge*. The sorted bam files were used to generate bed, bedgraph and bigwig formats for individual modifications.

**Peak-calling and analyses**

The reads were extracted from the bam to bed by the bedtools bamtobed option. Further reads were processed as mentioned in the SEACR (version 1.3) manual to get the bedgraph. These bedgraph files were subjected to peak calling through SEACR with a stringent p-value of ≤1e-6 with the norm and relaxed options.

Further bedtools with various options were used for transforming bed files, such as *intersect, closest, sample or shuffle*. GNU awk editor was used for processing the bed files wherever required. Chromatin state predictions for the histone modification peaks were performed using ChromHMM (Ernst and Kellis 2012) (v1.10). For further analyses, the reproducible peaks were obtained by performing an intersection between peak files for each CUT&Tag replicates for histone PTMs. Overlap between the peaks for histone modifications for the H9 cells was performed using the *Intervene* package. While the overlapping peak counts were plotted as Venn Diagram for each histone PTM and IgG, the peaks for histone PTM combinations were plotted as an *upset* plot showing the number of overlapping peaks (y-axis) along with the histone modification peak numbers on the x-axis.

**TE enrichment analyses**

Tracks for the repeats (rmsk) were obtained from the UCSC genome table browser for hg19 and hg38. Reproducible peaks (consistent between two replicates) were used to generate the Observed vs. expected frequency for different TE classes (*Alu*, full-length L1 and LTR), Gene-body and TSSs. These were calculated across various histone modification CUT&Tag peaks. The intersect count was obtained for each histone modification using bedtools (version v2.28.0) intersect (*bedtools intersect -wa -u options)* for the mentioned genomic elements. The expected occurrences in the genome were calculated by intersecting the genomic elements with the randomised genomic coordinates (number, length and chromosome id matched) across different histone modifications. The ratios for observed vs expected at these genomic elements for each histone PTM were calculated and used to plot as a heatmap using ggplot2 heatmap function in R.

**RepeatMasker**

To analyse the repeat content of the different histone modification peak sets, the fasta was obtained using the bedtools getfasta tool from hg38.fa reference genome. The sequences thus obtained were subjected to RepeatMasker (version 4.0.7) to get the repeat content across these genomic sequences for different histone modifications.

**Bigwig generation and plotting**

Sorted bam files were subjected to bigwig generation via deepTools (version 3.5.1) bamCoverage tool with *--binSize 20 –normalize Using BPM or CPM --scaleFactor=1.0 --smoothLength 60 --extendReads 150 --centerReads* options. The signal was normalised to IgG through bigwigCompare with option –operation first or subtract. The bigwig files were used for plotting signals or visualisation in the genome browser. The genome-browser views were obtained by viewing the signal tracks in the UCSC genome browser. For the knock-down (in H9 cells) or knock-out (in TDF cells) studies, the samples were normalised based on the number of reads mapping to the *E. coli* genome.

The plotting of signals at various genomic landmarks and bed coordinates was carried out through deepTools. Matrices were generated using deepTools *computeMatrix reference-point* or *scale-regions* option. These matrices were used for plotting heatmaps or average summary plots by *plotHeatmap* or *plotProfile* function in deepTools with or without clustering by the *k-means* algorithm. The sorted bam files were also used to study the correlation between the individual replicates for the CUT&Tag across histone PTMs and IgG using *multiBamSummary function in deeptools with options bins* and plotted as Pearson correlation heatmap using deeptools *plotCorrelation* function with options *--skipZeros.*

Further, H4K16ac signals from GSE84618 for brain (prefrontal lobe) tissues from Young, Alzheimer's Disease and Old individuals were compared for the TE elements. Similarly, the bigwigs were obtained for the proliferative and senescence model in IMR90 cells (GSM1358821) for L1 and LTR subfamilies. The signals were compared as heatmaps or average type summary profiles using above mentioned tools. The signals at the H4K16ac marked TEs were also plotted as average type summary plots using *plotProfile* function for transcription factors YY1 (ENCFF904SDR), RAD21 (ENCFF506AAX) and CTCF (ENCFF473IZV) using ENCODE datasets (bigwigs) normalized as fold change over control.

**TAD border annotation**

To call TADs in human embryonic stem cells (H9), we used Hi-C data for two replicates available from (Zhang et al., 2019) under accession numbers (GSM3262956 and GSM3262957). We first generated contact domains for all chromosomes at a 10 kb resolution using the Arrowhead tool from Juicer using Knight-Ruiz Normalisation (Durand et al., 2016). We extracted the borders of these TAD. To ensure we identify a robust set of TAD borders, we selected with a score above one that are common borders between the two replicates, assuming a maximum gap of 1 bin (10 kb). This resulted in 9952 robust TAD borders. The TAD calling was performed on the hg19 reference genome, and to allow integration with the rest of the analysis, we lifted over the common TAD borders from hg19 to hg38.

**Significant loops calling using Micro-C**

Chromatin loops were called with the HiCCUPS tool from the Juicer software suite (Durand et al., 2016) on micro-C data in H1 hESCs (Krietenstein et al., 2020). Loops were called using a 5 and 10 kb resolution, 10% FDR, Knight-Ruiz normalization, a window of 7 and 5, peak width of 2 4 and, thresholds for merging loops of 0.02, 1.5, 1.75, 2, and distance to merge peaks of 20 kb (--r 5000,10000 -k KR -f .1,.1 -p 4,2 -i 7,5 -t 0.02,1.5,1.75,2 -d 20000,20000).

**Motif enrichment analysis**

Enrichment of transcription factor binding sites (TFBS) at the TEs (>5kb L1, *Alu* and LTR) overlapping with histone modifications (H4K16ac, H3K27ac and H3K122ac) or a similar number of randomised genomic bins (chromosome, length matched) was performed. The experimentally determined TFBS for the H1-hESCs was fetched from the UCSC genome browser as TFBS clusters. The number of motifs for each TE class was either positive for histone modification or randomised genomic bins for histone modification for all TFBS. The internal distribution profile of motifs across each TEs was determined as percentage distribution and enrichment score defined as (Diff/Sum) of motif counts' percentage between observed (TEs positive for histone modification) vs expected (randomised genomic bins) occurrence of motifs. The ratios obtained for each TFBS were plotted as a heatmap using the R package *ComplexHeatmap.*

**ATAC-seq data analysis**

The ATAC-seq reads were processed for mapping by trimming for adapters using the Trimmomatic tool, followed by aligning to the hg38 genome through local Bowtie2(version) with these parameters for pair-end mapping: *--very-sensitive-local --no-unal --no-mixed --no-discordant --phred33 -I 10 -X 700 (Alteration in -X to 2000was done to allow the mapping of reads for H9 cells)*. The mapped reads were processed as described above (CUT&Tag data analysis) to generate the bigwigs. The signal was normalized as log2 fold changefor Control over MSL3 Knockdown using *bigwigCompare* function in deeptools with *--skipZeroOverZero --operation log2.* Using the same matrix generation and heatmap tools, further ATAC-seq signals were compared at the full-length L1 subfamilies and LTR sub-classes.

**RNA-seq data analysis**

The reads obtained from RNA-seq for H9 cells were mapped to the human genome using STAR following the Bluebee-CORALL pipeline of mapping. For TDFs, the RNA-seq datasets were downloaded from NCBI-GEO for accession ID GSE144019. The reads were mapped to hg38 following the same pipeline as H9 except for the single-end specification in TDFs.

For differential enrichment analysis, the fragment counts for each dataset were obtained using the *featurecounts* tool from the SubRead package. The GTF file for genes was obtained from ENSEMBL, and for different TE classes (*Alu*, L1 and LTR), it was fetched from the UCSC table browser. These feature counts were used for the differential enrichment analyses using the *DESeq2* package in R. The DESeq was performed with defaults. The differential expression of genes was visualised as a volcano plot. The differential gene expression table can be found in additional data.

The uniquely mapped reads were filtered using samtools for MAPQ of 255. Further, the unique alignments' bam files were merged, sorted and indexed using samtools, followed by bigwig generation using the deepTools function *bamCoverage*. The normalised signal was generated as log2 fold change for Control over MSL3 knockdown using *bigwigCompare* function in deeptools with *--operation log2*. The signal was compared at the full length L1 as well as genes.

The RNA signal across the various subfamilies of TEs as well as genes in the flanks (<10k, 10-25k and 25-50k) of the H4K16ac marked LTR and full length L1 were calculated as RPKM from the read counts obtained for each gene or TEs across the multiple replicates for H9 (Control or MSL3 knockdown) and TDF (WT or MSL1 knockout). The RPKM signal was then plotted as a violin and box-plot using *ggplot2 in R*. To compare, the same number of TEs (H4K16ac positive in TDFs) which are H4K16ac negative was obtained by subsampling using the *bedtools sample*. The signal was plotted as the *log10* value of the RPKM on the Y-axis. The RNA signals were plotted as violin plots with box plots with a median. The statistical analyses for all the violin plot comparisons were performed using *Dunn test* with *Bonferroni* correction.

**STARRseq data analysis**

To assess the potential of the TEs marked by H4K16ac to act as enhancers, we compared the STARR-seq signal in K562 (ENCFF611ZHY) and SH-SY5Y (ENCFF571ARG) cells at the TE elements: full-length L1 (>5kb) and ERV/LTRs. The signal was plotted as a heatmap from the start (L1) or centre (LTR) of the TE elements sorted according to the H4K16ac signal. Further, the signal was compared as violin plots for the four sets of peak combinations with respect to overlap among peaks that overlap with TEs. These were H3K4me1(+) only, H3K4me1(+)H3K27ac(+)H4K16ac(+), H4K16ac(+)H3K4me1(-) and H4K16ac(+)H3K4me1(+) peaks that overlap with LTRs.

**Statistical tests**

For all the RT-qPCRs, an *unpaired t-test with Welch correction* (two-stage step-up) was performed between the groups using *GraphPad Prism9*. For all the violin plots, the statistical tests were performed using the *dunn test* function in R tool *rstatix.* For multiple-group comparisons, Dunn's test with Bonferroni correction was used between the groups.

**Supplementary Figures**

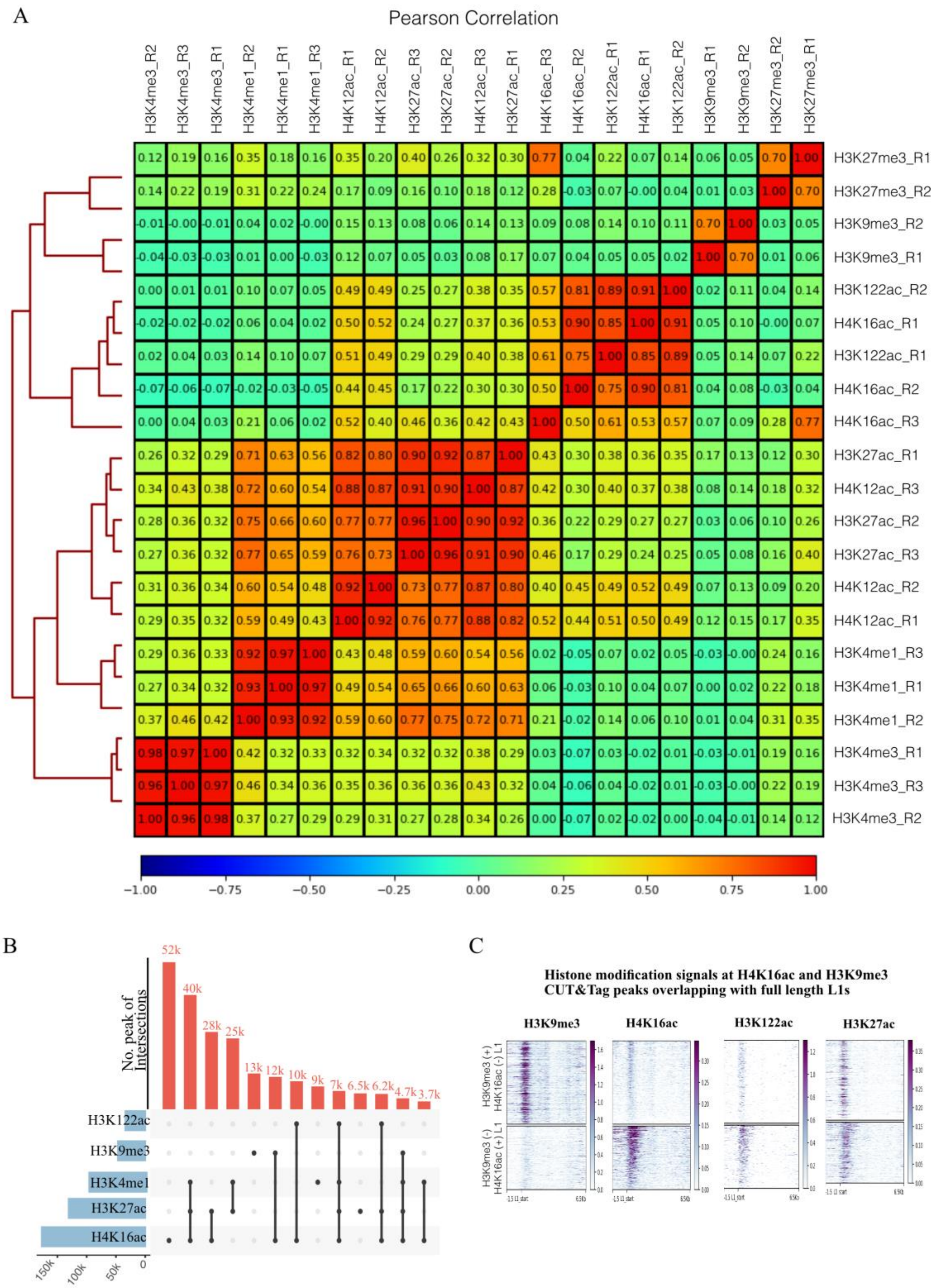

**Supplementary Fig. 1. Related to Fig. 1. CUT&Tag data showing enrichment of H4K16ac in TEs. A.** Pearson correlation heatmap for the CUT&Tag replicates across histone modifications in H9 cells. **B.** Upset plot showing the intersection of CUT&Tag peaks at TE (LTR, *Alu* and full-length L1) families. The X-axis shows the total number of peaks, and the Y-axis is the number of peaks intersected. **C.** Heatmaps showing signals (CPM) for the H3K9me3, H4K16ac, H3K27ac and H3K122ac at the full-length L1s marked by either H3K9me3 (top of each heatmap) or H4K16ac (bottom of each heatmap).

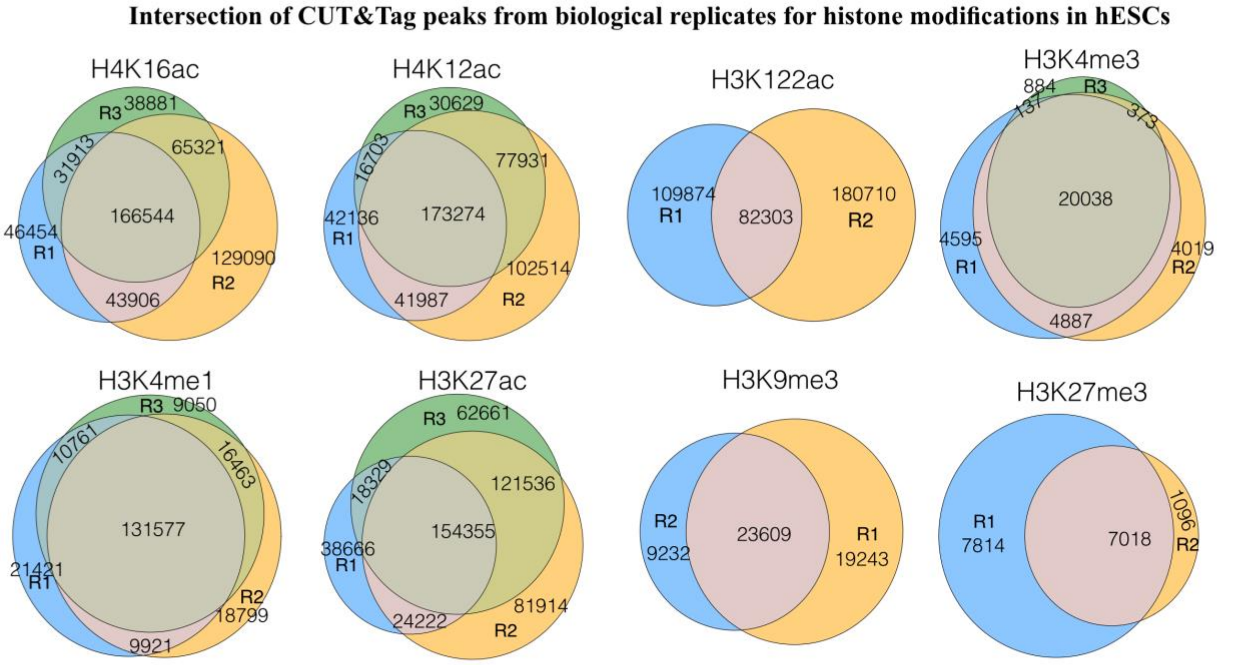

**Supplementary Fig. 2. Related to Fig. 1, H4K16ac is enriched at TEs in mESCs.**
**A.** Venn diagrams showing reproducibility for the CUT&Tag peaks among the replicates called for the Histone PTMs.

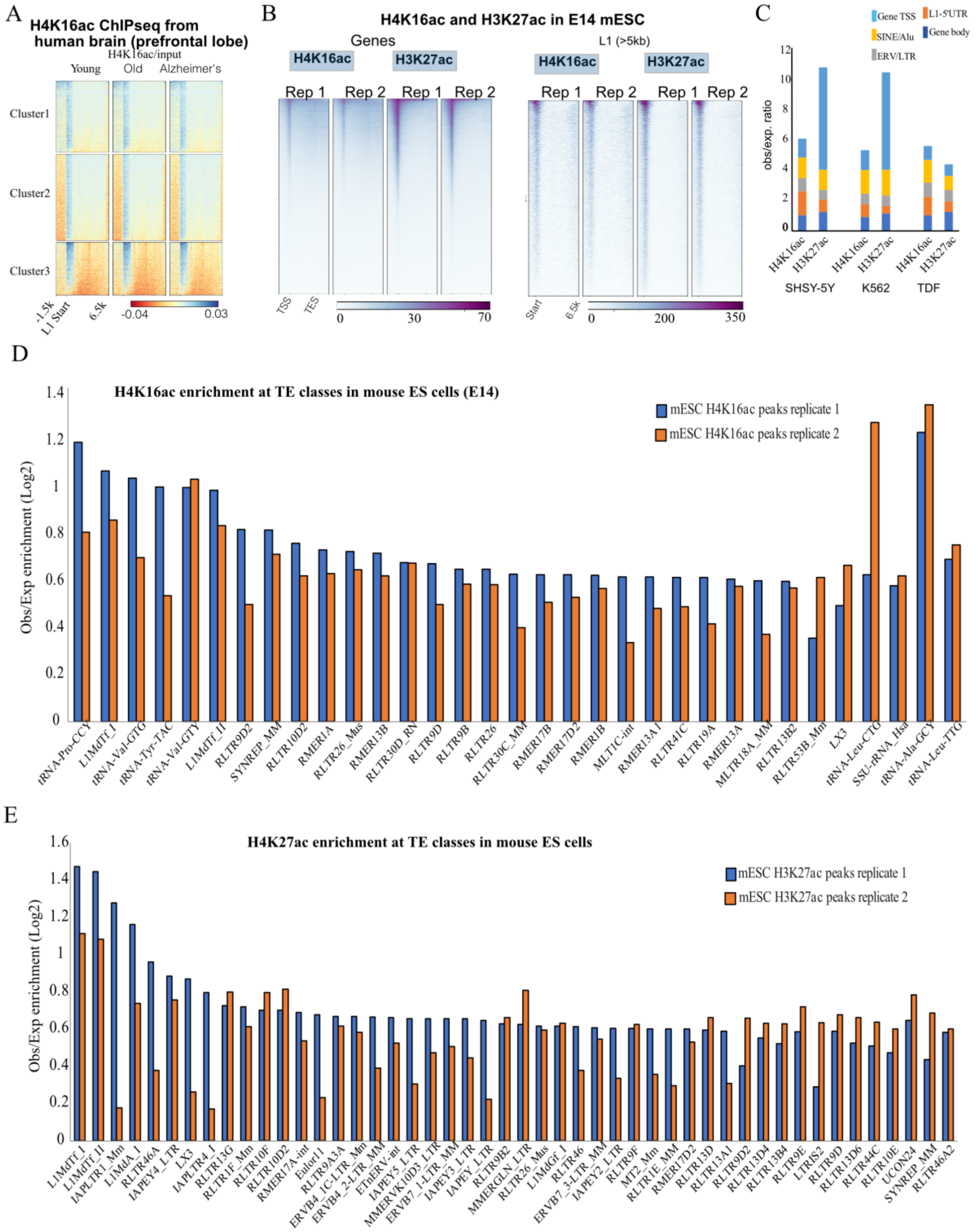

**Supplementary Fig. 3.** Related to Fig. 1 and 2. **A.** Heatmap showing H4K16ac ChIPseq signal across full-length L1s in human brain (prefrontal lobe) tissues from young, old and Alzheimer's patients from Nativio et al. 2018. **B.** Heatmap showing two replicates of H4K16ac and H3K27ac CUT&Tag signal across RefSeq genes (left) and full-length L1s (>5kb) from the mouse genome.

**C.** Stacked bar plot showing ratio (Y-axis) of observed over expected (background) for the TSS, genebody and TEs (LTR, Alu and L1) overlapping with H4K16ac or H3K27ac (X-axis) in SHSY-5Y, K562 and TDF cells. **D** and **E.** Observed over expected enrichment ratio for H4K16ac and H3K27ac mouse embryonic stem cells (E 14 mESCs) CUT&Tag peaks at transposable elements from mouse genome (from Repbase).

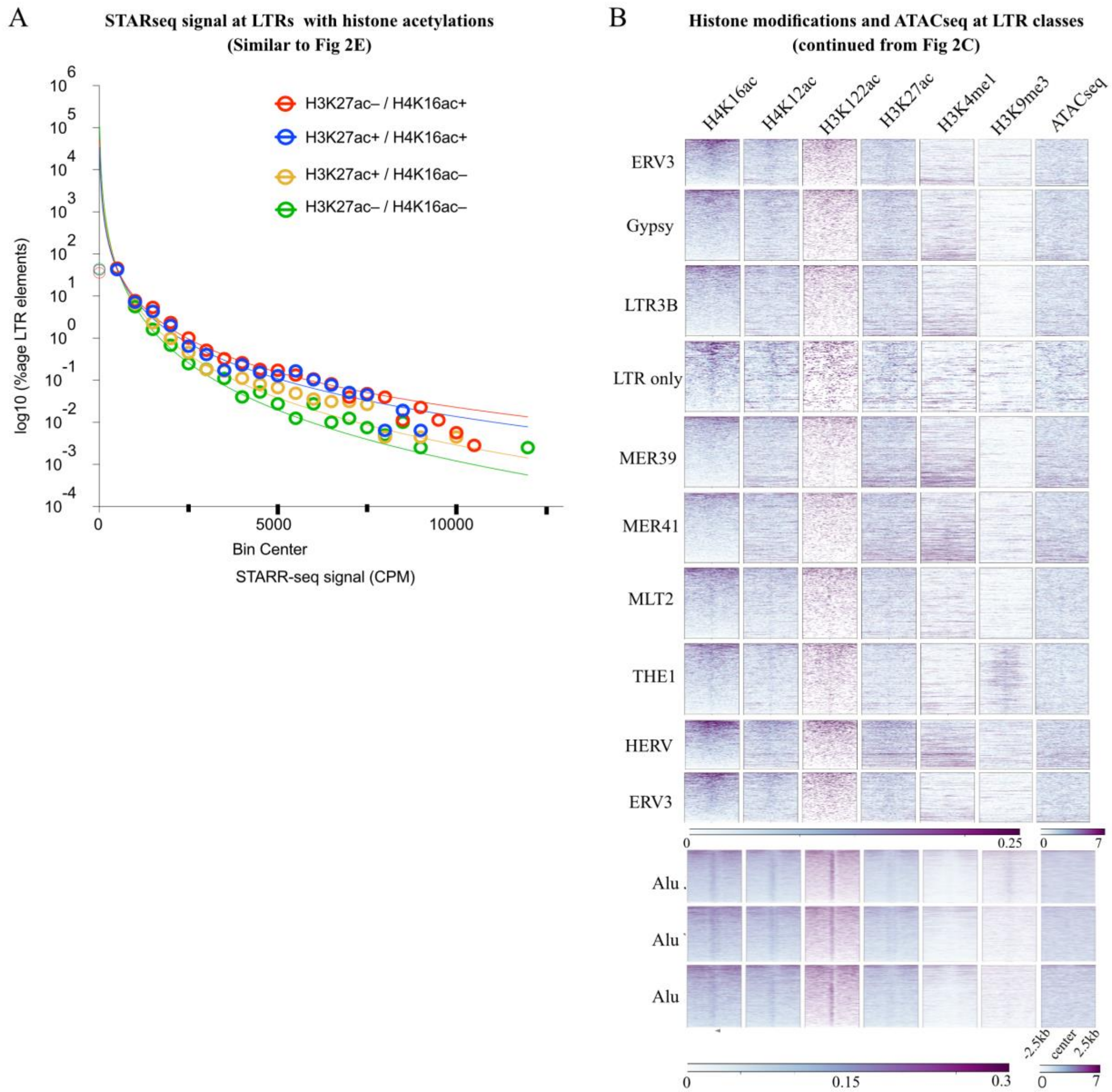

**Supplementary Fig. 4.** Related to Fig. 2. **A.** Frequency distribution of LTR elements (Y-axis, log 10 percentage) showing the STARR-seq signal enrichment (X-axis) that are H3K27ac-/H4K16ac+, H3K227ac+/H4K16ac+, H3K227ac+/H4K16ac- or H3K227ac-/H4K16ac-.
**B.** Heatmaps showing signals (CPM) for the H4K16ac, H4K12ac, H3K27ac and H3K122ac, H3K4me1, H3K9me3 and ATACseq at the LTR subfamilies and Alu subfamilies.

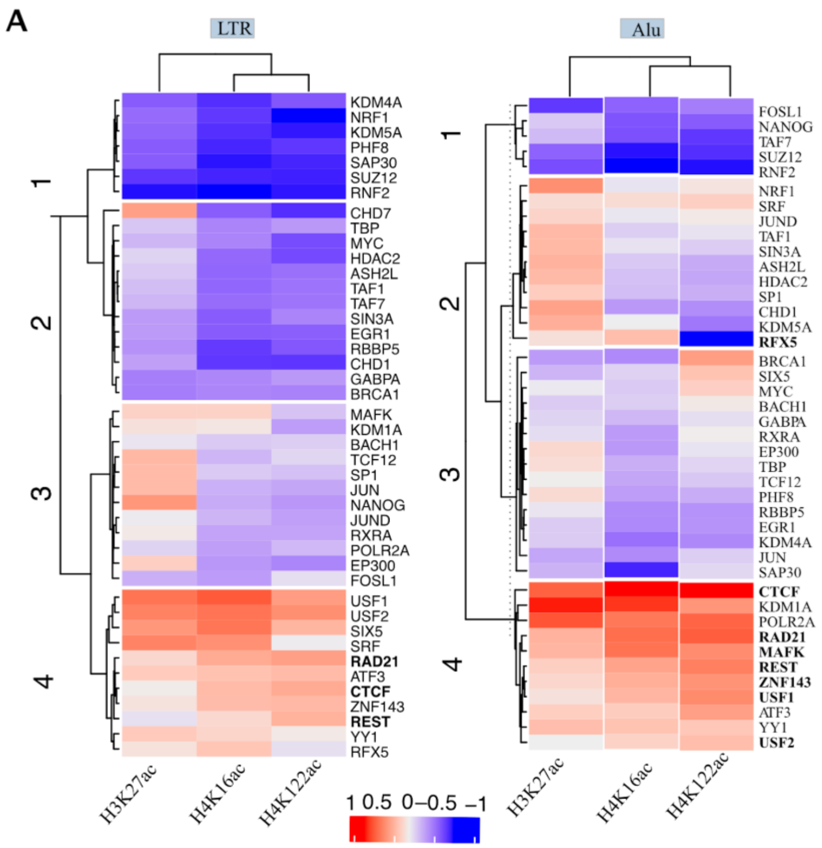

**Supplementary Fig. 5.** Related to Fig. 3. **A**) Like Fig. 3A, transcription factor binding sites enriched at the H3K27ac, H4K16ac and H3K122ac marked LTR and *Alu* in hESCs.

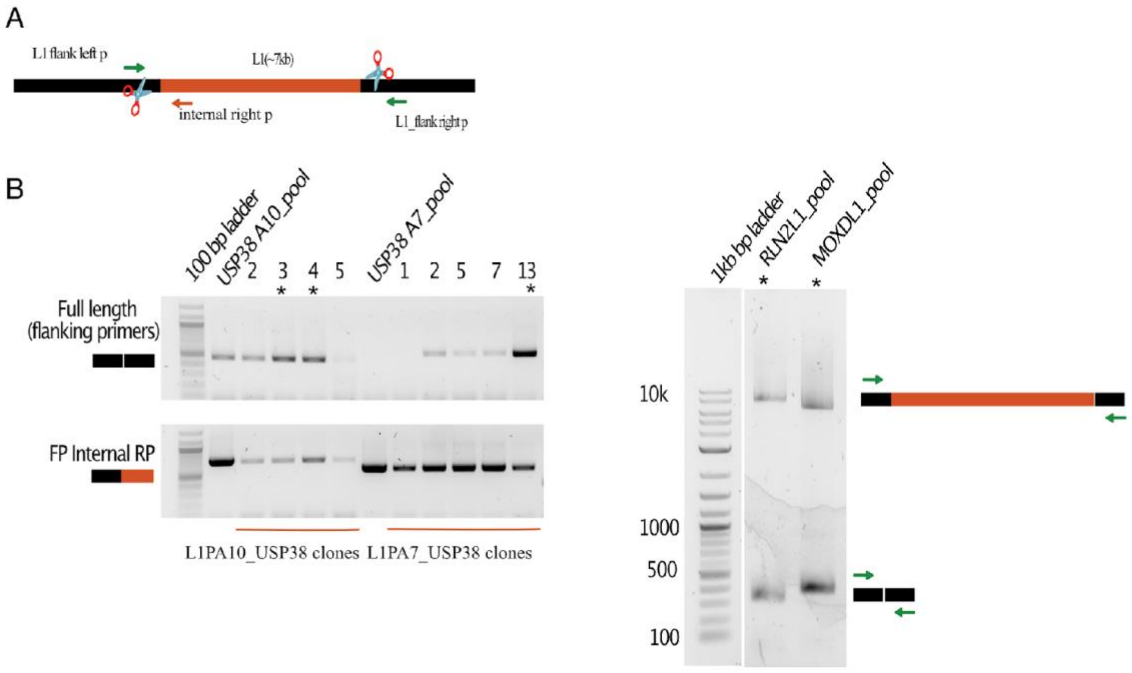

**Supplementary Fig. 6. Related to Fig 4. CRISPR CAS9 mediated deletion of L1 elements A)** Illustration showing the Full length LINE1 (L1, ~7kb), guideRNAs sites for CAS9 cutting (scissors), and the flanking primers (green arrow) and internal reverse primer (orange arrow) used for genotyping. B) Agarose gel electrophoresis showing PCR products for L1PA10 and L1PA7 clones, amplified using L1 flanking primers. ~500bp amplification showing deletion of L1 (above). PCR with internal reverse primers showing presence of wild type allele (below). C) PCR amplicons with L1 flanking primers for pool of cells showing nearly 50% deletion efficiency for L1PA7 at the RLN2 locus and L1PA8 at the MOXD1 locus.

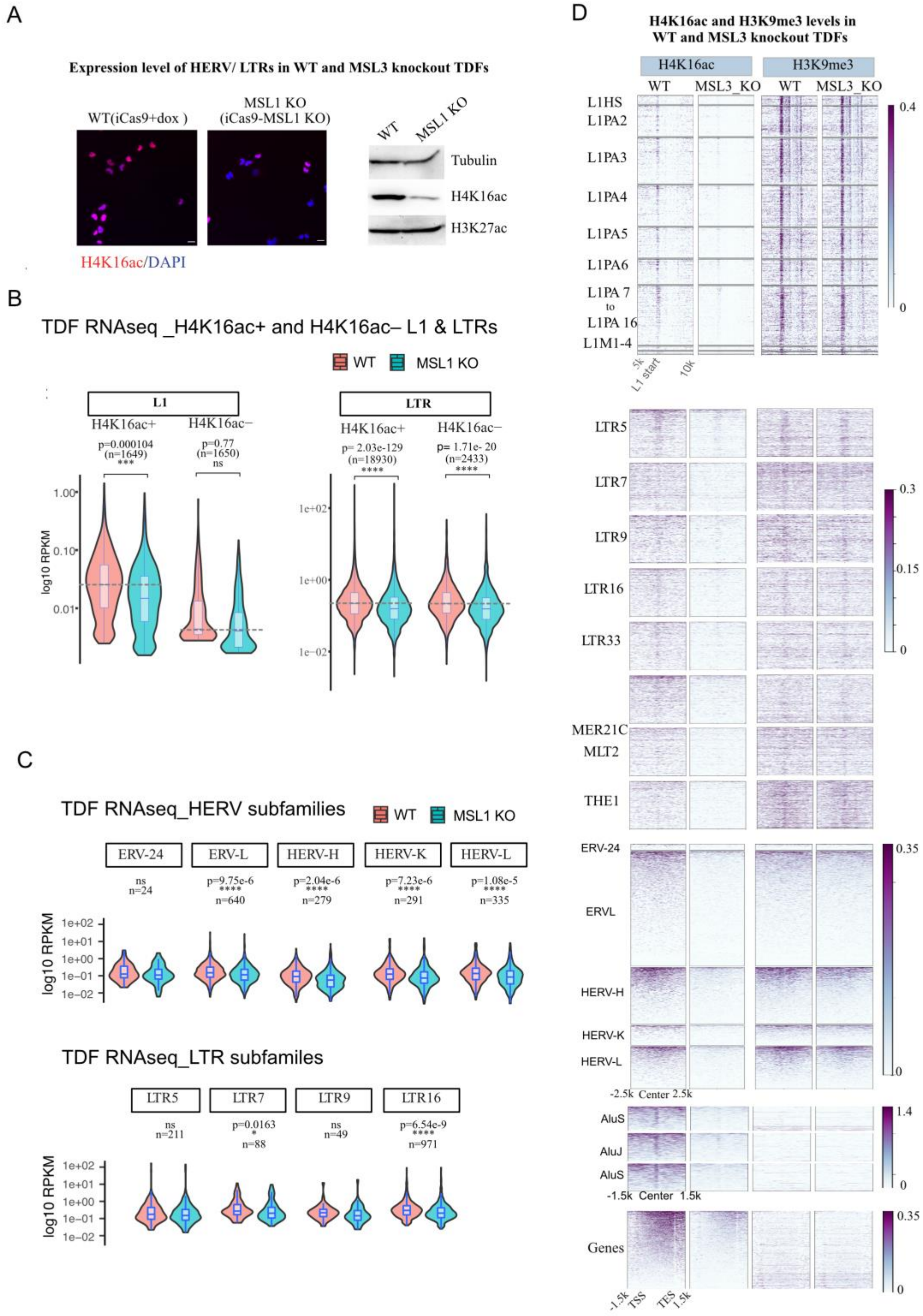

**Supplementary Fig. 7. Related to Fig. 5. Depletion of MSL proteins leads to downregulation of TEs A.** Immunofluorescence images showing H4K16ac levels (Magenta) in WT and MSL1 KO

TDFs (left). Western blots showing H4K16ac level in MSL1 KO and WT TDFs (Right). **B.** Violin plots showing the log10 RPKM signal of RNAseq reads for parental (WT) and doxycycline-inducible MSL1 (MSL1 KO) for L1s (left panel) and LTRs (right panel) that are either H4K16ac+ or H4K16ac-. **C.** Violin plots for RNAseq signal across different ERV subfamilies (ERV24, ERVL, HERVK, HERVH and HERVL; top), and LTR subfamilies (LTR5, LTR7, LTR9 and LTR16; below). Statistical tests for all violin plots were performed as Dunn test with Bonferroni correction. **D.** Heatmap comparing the CUT&Tag signals for H4K16ac and H3K9me3 for WT and MSL1 KO samples across L1 subfamilies, LTRs and ERV subfamilies, Alu subfamilies and NCBI refseq genes.

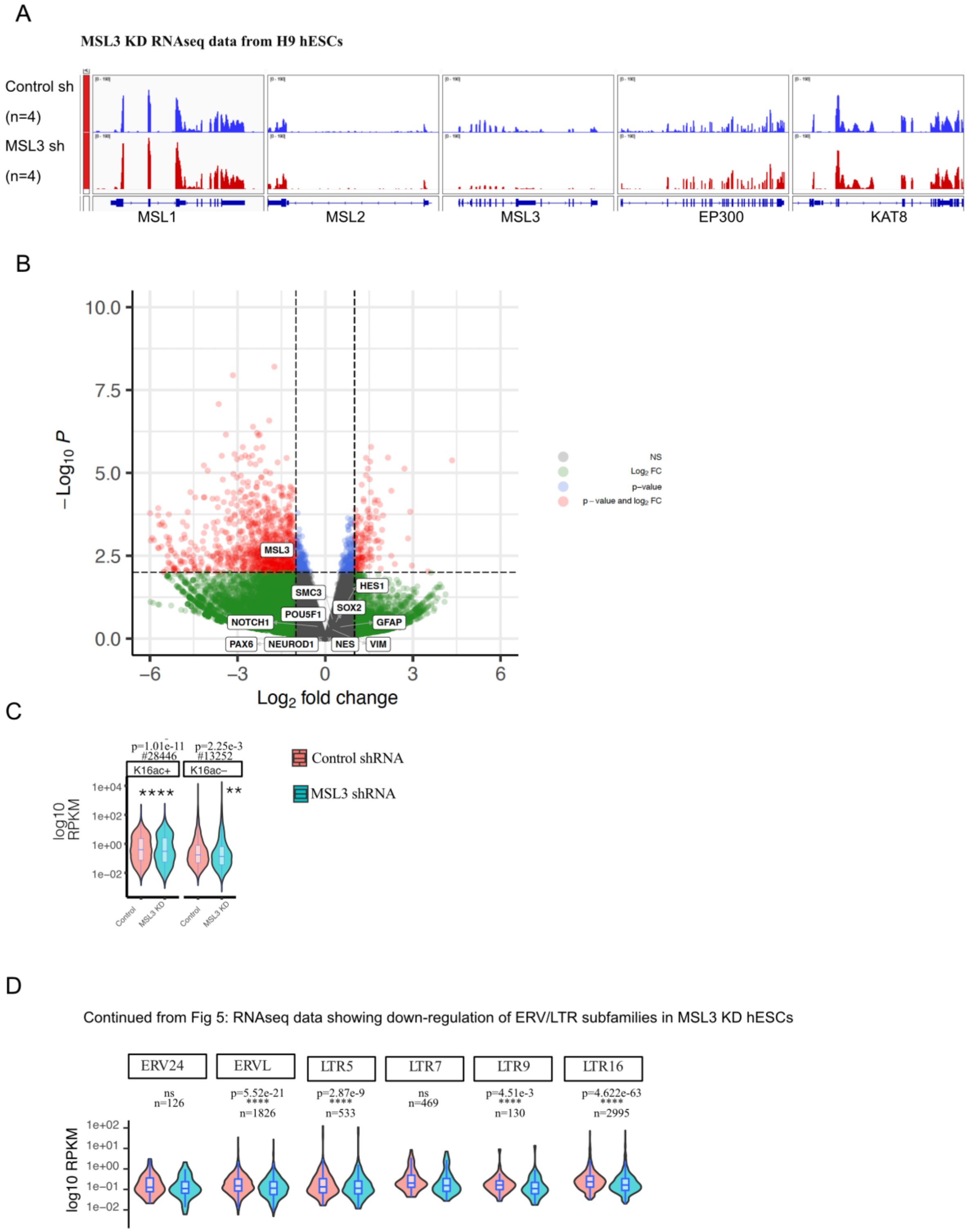
A
MSL3 KD RNAseq data from H9 hESCs
Control sh
(n=4)
MSL3 sh
(n=4)
MSL1
MSL2
MSL3
EP300
KAT8
B
10.0
7.5
5.0
2.5
0.0
−Log10 P
−6
−3
0
3
6
Log2 fold change
MSL3
SMC3
HES1
SOX2
POU5F1
NOTCH1
GFAP
PAX6
NEUROD1
NES
VIM
NS
Log2 FC
p−value
p − value and log2 FC
C
p=1.01e-11
#28446
p=2.25e-3
#13252
K16ac+
K16ac−
****
**
1e+04
1e+02
1e+00
1e−02
log10 RPKM
Control
MSL3 KD
Control
MSL3 KD
Control shRNA
MSL3 shRNA
D
Continued from Fig 5: RNAseq data showing down-regulation of ERV/LTR subfamilies in MSL3 KD hESCs
ERV24
ERVL
LTR5
LTR7
LTR9
LTR16
ns
n=126
p=5.52e-21
****
n=1826
p=2.87e-9
****
n=533
ns
n=469
p=4.51e-3
****
n=130
p=4.622e-63
****
n=2995
1e+02
1e+01
1e+00
1e−01
1e−02
log10 RPKM

**Supplementary Fig. 8. Related to Fig 5. A.** IGV browser tracks showing RNAseq reads (RPKM) at *MSL1*, *MSL2*, *MSL3* and *KAT8* locus in control knockdown (nontargeting shRNA) and MSL3 knockdown (n=4, biological replicates) H9 hESCs. **B.** Volcano plot showing up-and down-regulated genes upon lentiviral shRNA mediated knockdown of MSL3. Pluripotency-associated genes (e.g., POU5F1, NANOG, SOX2) and genes expressed in neuronal differentiation (e.g., PAX6, GFAP, NES, NEUROD1) are shown in arrow marks. **C.** Violin plots for the RNAseq signal (log10 RPKM) for the Control- and MSL3-shRNA for H9 ESCs at H4K16ac positive or H4K16ac negative genes. **D.** Like C but for LTR subfamilies (ERV24, ERVL, LTR5, LTR7, LTR9 and LTR16; bottom panel).

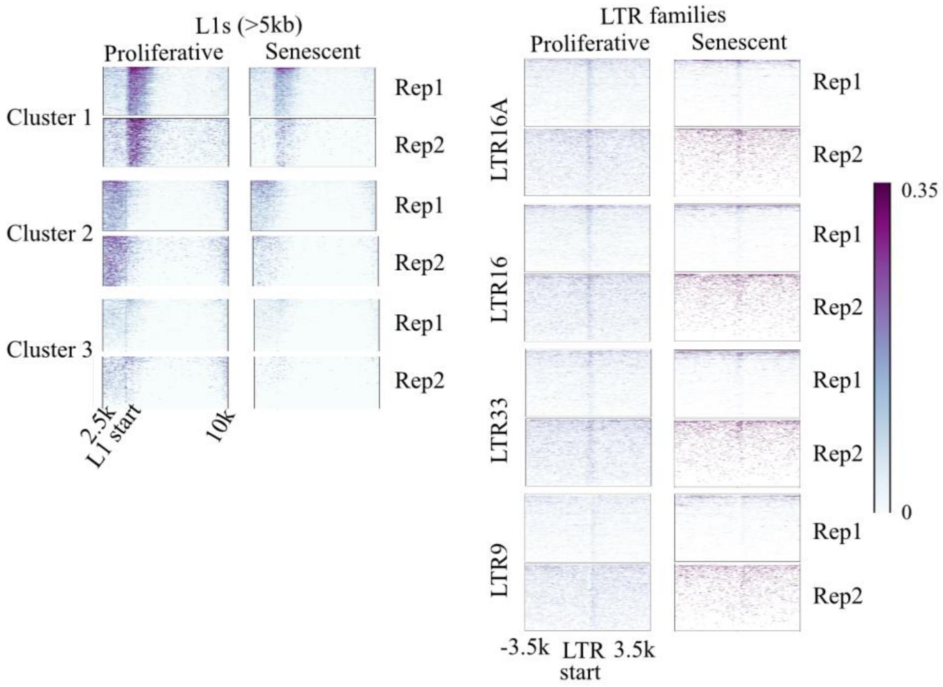

**Supplementary Fig. 9.** H4K16ac ChIPseq/input signal for L1 (left) and LTR (right) subfamilies in the proliferative and senescent IMR90 cell line.

**Supplementary Table 1. Data generated and analysed in this stud**

| | Experiment | GEO ID | Cell line, antibody, replicate number | Reference |
|---|---|---|---|---|
| 1 | CUT&Tag | GSM6043377 | H9, H3K27ac Rep1 | This study |
| 2 | CUT&Tag | GSM6043378 | H9, H3K27ac Rep2 | This study |
| 3 | CUT&Tag | GSM6043379 | H9, H3K27ac Rep3 | This study |
| 4 | CUT&Tag | GSM6043380 | H9, H3K27me3 Rep1 | This study |
| 5 | CUT&Tag | GSM6043381 | H9, H3K27me3 Rep2 | This study |
| 6 | CUT&Tag | GSM6043382 | H9, H3K36me3 Rep1 | This study |
| 7 | CUT&Tag | GSM6043383 | H9, H3K4me1 Rep1 | This study |
| 8 | CUT&Tag | GSM6043384 | H9, H3K4me1 Rep2 | This study |
| 9 | CUT&Tag | GSM6043385 | H9, H3K4me1 Rep3 | This study |
| 10 | CUT&Tag | GSM6043386 | H9, H3K4me3 Rep1 | This study |
| 11 | CUT&Tag | GSM6043387 | H9, H3K4me3 Rep2 | This study |
| 12 | CUT&Tag | GSM6043388 | H9, H3K4me3 Rep3 | This study |
| 13 | CUT&Tag | GSM6043389 | H9, H4K12ac Rep1 | This study |
| 14 | CUT&Tag | GSM6043390 | H9, H4K12ac Rep2 | This study |
| 15 | CUT&Tag | GSM6043391 | H9, H4K12ac Rep3 | This study |

| | | | | |
|---|---|---|---|---|
| 16 | CUT&Tag | GSM6043392 | H9, H4K16ac Rep1 | This study |
| 17 | CUT&Tag | GSM6043393 | H9, H4K16ac Rep2 | This study |
| 18 | CUT&Tag | GSM6043394 | H9, H4K16ac Rep3 | This study |
| 19 | CUT&Tag | GSM6043395 | H9, IgG Rep1 | This study |
| 20 | CUT&Tag | GSM6043396 | H9, IgG Rep2 | This study |
| 21 | CUT&Tag | GSM6043397 | H9, IgG Rep3 | This study |
| 22 | CUT&Tag | GSM6043398 | H9, H3K122ac Rep1 | This study |
| 23 | CUT&Tag | GSM6043399 | H9, H3K122ac Rep2 | This study |
| 24 | CUT&Tag | GSM6043400 | H9, H3K9me3 Rep1 | This study |
| 25 | CUT&Tag | GSM6043401 | H9, H3K9me3 Rep2 | This study |
| 26 | CUT&Tag | GSM6043402 | TDF, icas9 Igg | This study |
| 27 | CUT&Tag | GSM6043403 | TDF, icas9 H4K16ac Rep1 | This study |
| 28 | CUT&Tag | GSM6043404 | TDF, icas9 H3K27ac | This study |
| 29 | CUT&Tag | GSM6043405 | TDF, icas9_K9me3 | This study |
| 30 | CUT&Tag | GSM6043406 | TDF, MSL3KO_Igg | This study |
| 31 | CUT&Tag | GSM6043407 | TDF, MSL3KO H4K16ac | This study |
| 32 | CUT&Tag | GSM6043408 | TDF, MSL3KO H3K27ac | This study |
| 33 | CUT&Tag | GSM6043409 | TDF, MSL3KO H3K9me3 | This study |
| 34 | CUT&Tag | GSM6043410 | TDF, MSL1KO Igg | This study |
| 35 | CUT&Tag | GSM6043411 | TDF, icas9 H4K16ac Rep2 | This study |
| 36 | CUT&Tag | GSM6043412 | TDF, icas9 H4K16ac Rep3 | This study |
| 37 | CUT&Tag | GSM6043413 | TDF, MSL1KO H4K16ac Rep1 | This study |
| 38 | CUT&Tag | GSM6043414 | TDF, MSL1KO H4K16ac Rep2 | This study |
| 39 | CUT&Tag | GSM6043415 | HEK, IgG Rep1 | This study |

| | | | | |
|---|---|---|---|---|
| 40 | CUT&Tag | GSM6043416 | HEK, H4K16ac Rep1 | This study |
| 41 | CUT&Tag | GSM6043417 | HEK, H4K16ac Rep2 | This study |
| 42 | CUT&Tag | GSM6043418 | HEK, H3K27ac Rep1 | This study |
| 43 | CUT&Tag | GSM6043419 | HEK, H3K27ac Rep2 | This study |
| 44 | CUT&Tag | GSM6043420 | HeLa, IgG Rep1 | This study |
| 45 | CUT&Tag | GSM6043421 | HeLa, H4K16ac Rep1 | This study |
| 46 | CUT&Tag | GSM6043422 | HeLa, H4K16ac Rep2 | This study |
| 47 | CUT&Tag | GSM6043423 | HeLa, H3K27ac Rep1 | This study |
| 48 | CUT&Tag | GSM6043424 | HeLa, H3K27ac Rep2 | This study |
| 49 | CUT&Tag | GSM6043425 | K562, IgG Rep1 | This study |
| 50 | CUT&Tag | GSM6043426 | K562, H4K16ac Rep1 | This study |
| 51 | CUT&Tag | GSM6043427 | K562, H4K16ac Rep2 | This study |
| 52 | CUT&Tag | GSM6043428 | K562, H3K27ac Rep1 | This study |
| 53 | CUT&Tag | GSM6043429 | K562, H3K27ac Rep2 | This study |
| 54 | CUT&Tag | GSM6043430 | LNCaP, IgG Rep1 | This study |
| 55 | CUT&Tag | GSM6043431 | LNCaP, H4K16ac Rep1 | This study |
| 56 | CUT&Tag | GSM6043432 | LNCaP, H3K27ac Rep1 | This study |
| 57 | CUT&Tag | GSM6043433 | LNCaP, H3K27ac Rep2 | This study |
| 58 | CUT&Tag | GSM6043434 | PC3, IgG Rep1 | This study |
| 59 | CUT&Tag | GSM6043435 | PC3, H4K16ac Rep1 | This study |
| 60 | CUT&Tag | GSM6043436 | PC3, H4K16ac Rep2 | This study |
| 61 | CUT&Tag | GSM6043437 | PC3, H3K27ac Rep1 | This study |
| 62 | CUT&Tag | GSM6043438 | PC3, H3K27ac Rep2 | This study |
| 63 | CUT&Tag | GSM6043439 | RWPE, IgG Rep1 | This study |

| | | | | |
|---|---|---|---|---|
| 64 | CUT&Tag | GSM6043440 | RWPE, H4K16ac Rep1 | This study |
| 65 | CUT&Tag | GSM6043441 | RWPE, H3K27ac Rep1 | This study |
| 66 | CUT&Tag | GSM6043442 | SH-SY5Y, H4K16ac Rep1 | This study |
| 67 | CUT&Tag | GSM6043443 | SH-SY5Y, H4K16ac Rep2 | This study |
| 68 | CUT&Tag | GSM6043444 | SH-SY5Y, H3K27ac Rep1 | This study |
| 69 | CUT&Tag | GSM6043445 | SH-SY5Y, H3K27ac Rep2 | This study |
| 70 | ATAC-seq | GSM6043367 | H9, Scr ATAC Rep1 | This study |
| 71 | ATAC-seq | GSM6043368 | H9, Scr ATAC Rep2 | This study |
| 72 | ATAC-seq | GSM6043369 | H9, Scr ATAC Rep3 | This study |
| 73 | ATAC-seq | GSM6043370 | H9, MSL3 KD ATAC Rep1 | This study |
| 74 | ATAC-seq | GSM6043371 | H9, MSL3 KD ATAC Rep2 | This study |
| 75 | ATAC-seq | GSM6043372 | H9, MSL3 KD ATAC Rep3 | This study |
| 76 | ATAC-seq | GSM6043373 | TDF, icas9 ATAC Rep1 | This study |
| 77 | ATAC-seq | GSM6043374 | TDF, icas9 ATAC Rep2 | This study |
| 78 | ATAC-seq | GSM6043375 | TDF, MSL1 KO ATAC Rep1 | This study |
| 79 | ATAC-seq | GSM6043376 | TDF, MSL1 KO ATAC Rep2 | This study |
| 80 | ATACseq | GSM4770996 | THP1, sgNegCtrl ATAC | (Radzisheuskaya et al., 2021) |
| 81 | ATACseq | GSM4770997 | THP1, sgMSL1_KO ATAC | (Radzisheuskaya et al., 2021) |
| 82 | RNA-seq | GSM6043446 | H9, Scr RNA-seq Rep1 | This study |
| 83 | RNA-seq | GSM6043447 | H9, Scr RNA-seq Rep2 | This study |

| | | | | |
|---|---|---|---|---|
| 84 | RNA-seq | GSM6043448 | H9, MSL3KD RNA-seq Rep1 | This study |
| 85 | RNA-seq | GSM6043449 | H9, MSL3KD RNA-seq Rep2 | This study |
| 94 | Hi-C | GSM3262956 | H9, HiC.Rep1 | (Zhang et al., 2019) |
| 95 | Hi-C | GSM3262957 | H9, HiC.Rep2 | (Zhang et al., 2019) |
| 96 | ChIP-seq | GSM1358821 | IMR90, H4K16ac Prolif ChIP Rep1 | (Rai et al., 2014) |
| 97 | ChIP-seq | GSM1358822 | IMR90, H4K16ac Prolif ChIP Rep2 | (Rai et al., 2014) |
| 98 | ChIP-seq | GSM1358823 | IMR90, H4K16ac Senes ChIP Rep 1 | (Rai et al., 2014) |
| 99 | ChIP-seq | GSM1358824 | IMR90, H4K16ac Senes ChIP Rep 2 | (Rai et al., 2014) |
| 100 | ChIPseq | GSE84618 | Brain, H4K16ac-Input.AD.bw | (Nativio et al., 2018) |
| 101 | ChIPseq | GSE84618 | Brain, H4K16ac-Input.Old.bw | (Nativio et al., 2018) |
| 102 | ChIPseq | GSE84618 | Brain, H4K16ac-Input.Young.bw | (Nativio et al., 2018) |
| 103 | RNA-seq | GSM4278153 | TDF, WT Rep 1 | (Monserrat et al., 2021) |
| 104 | RNA-seq | GSM4278154 | TDF, WT Rep 2 | (Monserrat et al., 2021) |
| 105 | RNA-seq | GSM4278155 | TDF, WT Rep 3 | |

| | | | | |
|---|---|---|---|---|
| 106 | RNA-seq | GSM4278141 | TDF, MSL1 KO Rep 1 | |
| 106 | RNA-seq | GSM4278141 | TDF, MSL1 KO Rep 1 | |
| 107 | RNA-seq | GSM4278142 | TDF, MSL1 KO Rep 2 | |
| 108 | RNA-seq | GSM4278143 | TDF, MSL1 KO Rep 3 | |
| 109 | RNA-seq | GSM4278147 | TDF, MSL3 KO Rep 1 | |
| 110 | RNA-seq | GSM4278148 | TDF, MSL3 KO Rep 2 | |
| 111 | RNA-seq | GSM4278149 | TDF, MSL3 KO Rep 3 | |
| 112 | ATAC-seq | GSM5219560 | H9 hESC WT_ATAC-seq Rep1 | (Hsieh et al., 2022) |
| 113 | ATAC-seq | GSM5219561 | H9 hESC WT_ATAC-seq Rep2 | (Hsieh et al., 2022) |

Supplementary Table 2
Mapping details and number of peaks detected in all replicates across histone modifications

| Sample | Total read pairs (post trimming) | #unique aligned | #multi-aligned | unique aligned %age | multi-aligned %age | # of peaks |
|---|---|---|---|---|---|---|
| H9, H3K27ac Rep1 | 5267795 | 3461853 | 1177926 | 65.72 | 22.36 | 235572 |
| H9, H3K27ac Rep2 | 24283828 | 17273549 | 5108095 | 71.13 | 21.03 | 332822 |
| H9, H3K27ac Rep3 | 21041964 | 14397784 | 5081506 | 68.42 | 24.15 | 317466 |
| H9, H3K27me3 Rep1 | 1059518 | 512357 | 158594 | 48.36 | 14.97 | 14832 |
| H9, H3K27me3 Rep2 | 2330065 | 1682644 | 408995 | 72.21 | 17.55 | 4921 |
| H9, H3K4me1 Rep1 | 8726469 | 6222892 | 1431697 | 71.31 | 16.41 | 173680 |
| H9, H3K4me1 Rep2 | 16756188 | 12116934 | 3139439 | 72.31 | 18.74 | 158201 |
| H9, H3K4me1 Rep3 | 35390485 | 27655989 | 6124551 | 78.15 | 17.31 | 141821 |
| H9, H3K4me3 Rep1 | 6822949 | 5261704 | 791453 | 77.12 | 11.60 | 29657 |
| H9, H3K4me3 Rep2 | 40759484 | 33771216 | 4452970 | 82.85 | 10.92 | 28240 |
| H9, H3K4me3 Rep3 | 1452386 | 1152155 | 183540 | 79.33 | 12.64 | 24204 |
| H9, H4K12ac Rep1 | 7805697 | 4827628 | 1887568 | 61.85 | 24.18 | 274100 |
| H9, H4K12ac Rep2 | 23829711 | 15960547 | 5831918 | 66.98 | 24.47 | 345073 |
| H9, H4K12ac Rep3 | 27318999 | 17788703 | 8024831 | 65.11 | 29.37 | 253197 |
| H9, H4K16ac Rep1 | 8831804 | 5604427 | 2477020 | 63.46 | 28.05 | 288817 |
| H9, H4K16ac Rep2 | 18212861 | 10648064 | 4116399 | 58.46 | 22.60 | 370833 |

| | | | | | | |
|---|---|---|---|---|---|---|
| H9, H4K16ac Rep3 | 21797848 | 13885613 | 6910663 | 63.70 | 31.70 | 256460 |
| H9, H3K122ac Rep1 | 1434855 | 733254 | 532893 | 51.10 | 37.14 | 192177 |
| H9, H3K122ac Rep2 | 2802275 | 1621322 | 1077241 | 57.86 | 38.44 | 251153 |
| H9, H3K9me3 Rep1 | 16787273 | 6343184 | 10151605 | 37.79 | 60.47 | 43032 |
| H9, H3K9me3 Rep2 | 1776891 | 380612 | 1239764 | 21.42 | 69.77 | 43921 |
| H9, IgG Rep1 | 1455306 | 509592 | 397818 | 35.02 | 27.34 | 11089 |
| H9, IgG Rep2 | 1000089 | 386107 | 316372 | 38.61 | 31.63 | 10193 |
| H9, IgG Rep3 | 10973836 | 6075158 | 3832333 | 55.36 | 34.92 | 60305 |

Supplementary Table 3
Statistics for the histone modifications' reproducible peaks at genomic elements

| A) Ratio of enrichment over background for genomic elements overlapping histone modification peaks | | | | | |
|---|---|---|---|---|---|
| | Alu | L1 | LTR | TSS | Genes |
| IgG | 0.69 | 0.60 | 0.32 | 25.35 | 2.34 |
| H3K122ac | 0.60 | **5.45** | 0.76 | **2.58** | 1.09 |
| H3K12ac | **2.70** | 2.60 | 1.40 | 15.60 | 2.04 |
| H4K16ac | **1.89** | **2.93** | **1.86** | **1.02** | 1.31 |
| H3K27ac | **2.21** | 1.54 | 1.58 | 12.22 | 1.80 |
| H3K27me3 | 0.48 | 0.08 | 0.32 | 11.83 | 1.68 |
| H3K4me1 | 1.14 | 1.67 | 1.17 | 16.68 | 1.71 |
| H3K4me3 | 0.86 | 1.57 | 0.39 | 63.32 | 2.78 |
| H3K9me3 | 0.65 | **19.18** | **1.98** | **0.46** | 0.67 |
| B) Percentage of genomic elements overlapping reproducible peaks of histone modifications | | | | | |
| | %age of Alu Elements | %age of L1 Elements | %age of LTR elements | %age of TSSs | %age of Genes |

| **# of Genomic Elements** | **1204532** | **10538** | **764056** | **173733** | **38956** |
|---|---|---|---|---|---|
| IgG | 0.73 | 0.45 | 0.27 | 19.26 | 24.66 |
| H3K122ac | 1.5 | 15.83 | 1.85 | 4.39 | 41.44 |
| H3K12ac | **42.92** | 50.63 | 29.77 | **76.5** | 80.53 |
| H4K16ac | **35.76** | **51.47** | **34.89** | 22.86 | 67.99 |
| H3K27ac | **39.63** | 40.15 | **33.43** | **74.44** | 79.03 |
| H3K27me3 | 0.36 | 0.06 | 0.21 | 6.76 | 10.14 |
| H3K4me1 | 17.88 | 29.11 | 18.23 | **71.55** | 75.25 |
| H3K4me3 | 2.86 | 5.1 | 1.12 | **60.68** | 60.67 |
| H3K9me3 | 2.89 | **47.68** | 8.05 | 1.87 | 15.78 |

C) Percentage of reproducible peaks overlapping genomic elements

| | **Total reproducible peaks** | %age of Peaks at Alu | %age of Peaks at L1 | %age of Peaks at LTR | %age of Peaks at TSS | %age of Peaks at Genes |
|---|---|---|---|---|---|---|
| IgG | **7966** | 48.78 | 0.59 | 15.15 | 59.49 | 80.14 |
| H3K122ac | **69622** | 19.43 | 2.44 | 17.31 | 3.35 | 66.65 |
| H3K12ac | **220291** | 64.74 | 2.42 | 45.21 | 11.85 | 57.46 |
| H4K16ac | **217029** | 61.88 | 2.5 | 50.4 | 6.19 | 50.17 |
| H3K27ac | **229431** | 63.11 | 1.84 | 46.98 | 11.39 | 54.95 |
| H3K27me3 | **3820** | 42.93 | 0.16 | 18.82 | 62.28 | 74.08 |
| H3K4me1 | **132905** | 58.71 | 2.31 | 44.87 | 16.89 | 56.37 |
| H3K4me3 | **24013** | 56.7 | 2.24 | 20.04 | 64.53 | 77.9 |
| H3K9me3 | **23612** | 54.39 | 21.03 | 67.63 | 5.38 | 33.86 |

**Supplementary Table 4. Oligos used for RT-qPCR**

| | | |
|---|---|---|
| B-ACTIN | CAGCCATGTACGTTGCTATCCAGG | AGGTCCAGACGCAGGAT GGCATG |
| L1HS | CAAACACCGCATATTCTCACTCA | TGTGTCATCTAGCATTAGGTAT |
| L1PB | GAT GTT GGC GTG GAT GTG GT | TGG GAT TGC TGG ATC AAA TGG T |
| L1M | GGG GAG GTG GGG ATG GTT AA | GGT ATC CAT CCC CTC AAG CAT |
| L1PA 15/16 | CCATTTGACCCAGCAATCCCA | CCTCCAGCTGCATCCATGTT |
| HERVK gag | AGC AGG TCA GGT GCC TGT AAC ATT | TGG TGC CGT AGG ATT AAG TCT CCT |
| HERVH gag | CTT TTA TTA CCC AAT CTG CTC CCG AYA T | TTT AGT GGT GGA CAG TCT CTT TTC CAR TG |
| MSL3 | GTTATGCCACATGCCAACAT | CCAACGGGAGAGAGTGTAATCAA |
| NUS | TTCGGTCCTGTGGACAGCAC | CAGACGCTGTTCACAGGCTG |
| PEX1 | ATCAGAACTTGGAAATGGAACCT | CCTGAGTCATGGAGCTTGGT |
| GATAD1 | TCCACCAAAGGAAAAGGGAG | TGGAAACTGACTCAGGAGCTTT |
| USP38 | CACTCCTGAAAGGACTGGCA | AACTGCAAGAGCACCAGGTC |
| TANC2 | AAGATCCCAGAGAGAAATTTGGA | AGCAAGGTCTTCCAAGTGGG |
| MOXD1 | TCATTGGGGTTAAGGAGATCTACAG | AGCATCCATGAAAGAAAGGTTTT |
| COMMD10 | CAGTTGAAAAGTTCCGGCAG | TGAAGGTTAAGCTGCCATCC |

| | | |
|---|---|---|
| CYB561 | TGGTCATAGGCCTGATCTTCC | GACCTTGGTGGTGCGTTTAG |
| GAB1 | GGATGTCGCCTTCACGTAGT | ACTTGGAAATGCTCGTGGAA |
| PLGRKT | GATTGCGTGGTCTCGGGAATTC | AACAATCGGGACCAGGAAGGCT |
| ENPP1 | TGCACAGATGCCTGAACCT | ACATCCCCCAAATATTTATTCAGAT |
| STX7 | GGAATGATGATTCATGAACAAGG | CCCTTGACAGCTGCTGATTT |
| RLN2 | GATGCTCCTCAGACACCTAGAC | GTTGTAGCTGTGGTAATGCTGGC |
| SMARCA5 | CCTTTGAAGATGAAACCAGGG | GCTCTGTTCTACGGTGTCGGT |
| | **Genotyping primers for CRISPR deletions** | |
| L1PA10_USP38 | GGTGTGAGTGTGAATGAAACAG | GGAGCTAGGTGAAATGTACACA |
| L1PA7_USP38 | AATATCCTACAAGAGCAGTATGGTGC | GGGGCTCAAAATTAGCAAGG |
| L1PA7_RLN2 | CATAAGGAGGAAGGCCTCTATGC | ACTGCTTCTGATGGTATGTCCG |
| L1PA8_MOXD1 | CCACTGCAGTGTATTAAGAGGTG | TCCAAGGCTCAGGAAATCTG |

**Supplementary Table 5. Details of TEs used for CRISPRi and guide RNAs**

| GuideRNA name | gRNA (crRNA) sequence | putative target gene |
|---|---|---|
| L1PA10 g1 | ATGACTGAGTTTCTGAGGGG | USP38 |
| L1PA10 g2 | GGGTGGAGCAATGGCCTACT | |
| L1PA10 g1 | CCGCAGTCTCGGTTGATCAG | TANC2 |
| L1PA10 g2 | CTGGAAACTGCCTAAGACTA | |
| L1PA8g1 | CCTGGGAAGTGGCTAGTCTG | MOXD1 |
| L1PA8g2 | CAGGAGTTCTTACACATCAC | |
| L1PA7g1 | CCTCTCGGTACAGTCTCTTA | COMMD10 |
| L1PA7g2 | TAGCAGCGAGAATTTCACGT | |
| HERVH_g1 | TCAAGCTCGGGTCAAGCTCG | NUS1 |
| HERVH_g2 | ATACTTGTGGATTTAAGGTG | |
| HERVH_g1 | ACGAGTTGGGTGCTACAGGG | PEX1/GATAD1 |
| HERVH_g2 | AAGTGGTTGATTATACTGGG | |
| USP38L1PA2_noAc_g1 | CATGTTGGCAGTATGAAGTG | USP38 |
| USP38L1PA2_noAc_g2 | AACTGATCTAATTTAGAGGG | |
| USP38L1MA2_noAc_g1 | GGCAATAGATCATAGAGTCT | USP38 |
| USP38L1MA 2_noAc_g2 | TGACTTATATATCTAGGTGC | |
| L1PA10-USP38 gRNA UP | TACCACAGCTGTTTCTGGTC | USP38 |
| L1PA10-USP38 gRNA2 Down | TAACCCAAATAGTTATCTAC | |
| L1PA7-USP38 gRNA UP | CTTGAAGCAACAATCTGGTA | USP38 |
| L1PA7-USP38 gRNA2 Down | CATCAAGTCCTCTAGATATT | |

| | | |
|---|---|---|
| L1PA7-MOXD1gRNA UP | AGCCAACTAAGAGACTCCCC | MOXD1 |
| L1PA7-MOXD1gRNA Down | TCGTCCAGAGAGATCTCCCA | |
| L1PA7- RLN2gRNA up | CAGATACTTACTTACTCCAT | RLN2 |
| L1PA7-RLN2gRNA2 down | GTTACTACGTATGGTTAAAG | |

**Supplementary Table 6 Software and bioinformatic tools used in the study**

| RepeatMasker | https://www.repeatmasker.org | Reference |
|---|---|---|
| Bowtie2 | http://bowtie-bio.sourceforge.net/bowtie2/ | (Langmead and Salzberg, 2012) |
| STAR | https://github.com/alexdobin/STAR | (Dobin et al., 2013) |
| SAMtools | http://www.htslib.org/ | (Danecek et al., 2021) |
| BEDTools | https://github.com/arq5x/bedtools2/ | (Quinlan and Hall, 2010) |
| deepTools | https://github.com/deeptools/deepTools | (Ramírez et al., 2014) |
| ChromHMM | http://compbio.mit.edu/ChromHMM/ | (Ernst and Kellis, 2012) |
| ComplexHeatmap | https://jokergoo.github.io/ComplexHeatmap/ | (Gu et al., 2016) |
| DESeq2 | https://github.com/mikelove/DESeq2/ | (Love et al., 2014) |
| Intervene | https://github.com/asntech/intervene | (Khan and Mathelier, 2017) |
| SEACR | https://github.com/FredHutch/SEACR | (Meers et al., 2019) |

Durand, N.C., Shamim, M.S., Machol, I., Rao, S.S.P., Huntley, M.H., Lander, E.S., Aiden, E.L.,

Durand, N.C., Shamim, M.S., Machol, I., et al. (2016). Juicer Provides a One-Click System for Analyzing Tool Juicer Provides a One-Click System for Analyzing Loop-Resolution Hi-C Experiments. Cell Syst. *3*, 95–98.
Ernst, J., and Kellis, M. (2012). ChromHMM: Automating chromatin-state discovery and characterization. Nat. Methods *9*, 215–216.
Gu, Z., Eils, R., and Schlesner, M. (2016). Complex heatmaps reveal patterns and correlations in multidimensional genomic data. Bioinformatics *32*, 2847–2849.
Hsieh, F.K., Ji, F., Damle, M., Sadreyev, R.I., and Kingston, R.E. (2022). HERVH-derived lncRNAs negatively regulate chromatin targeting and remodeling mediated by CHD7. Life Sci. Alliance *5*, 1–19.
Kaya-okur, H.S., Wu, S.J., Codomo, C.A., Pledger, E.S., Bryson, T.D., Henikoff, J.G., Ahmad, K., and Henikoff, S. (2019). CUT&Tag for ef fi cient epigenomic pro fi ling of small samples and single cells. Nat. Commun. *10*, 1–10.
Khan, A., and Mathelier, A. (2017). Intervene: A tool for intersection and visualization of multiple gene or genomic region sets. BMC Bioinformatics *18*, 1–8.
Krietenstein, N., Abraham, S., Venev, S. V., Abdennur, N., Gibcus, J., Hsieh, T.H.S., Parsi, K.M., Yang, L., Maehr, R., Mirny, L.A., et al. (2020). Ultrastructural Details of Mammalian Chromosome Architecture. Mol. Cell *78*, 554-565.e7.
Langmead, B., and Salzberg, S.L. (2012). Fast gapped-read alignment with Bowtie 2. Nat. Methods *9*, 357–359.
Love, M.I., Huber, W., and Anders, S. (2014). Moderated estimation of fold change and dispersion for RNA-seq data with DESeq2. Genome Biol. *15*, 1–21.
Meers, M.P., Tenenbaum, D., and Henikoff, S. (2019). Peak calling by Sparse Enrichment Analysis for CUT & RUN chromatin profiling. Epigenetics Chromatin 1–11.
Monserrat, J., Morales Torres, C., Richardson, L., Wilson, T.S., Patel, H., Domart, M.C., Horswell, S., Song, O.R., Jiang, M., Crawford, M., et al. (2021). Disruption of the MSL complex inhibits tumour maintenance by exacerbating chromosomal instability. Nat. Cell Biol. *23*, 401–412.
Nativio, R., Donahue, G., Berson, A., Lan, Y., Amlie-Wolf, A., Tuzer, F., Toledo, J.B., Gosai, S.J., Gregory, B.D., Torres, C., et al. (2018). Publisher Correction: Dysregulation of the epigenetic landscape of normal aging in Alzheimer’s disease (Nature Neuroscience, (2018), 21, 4, (497-505), 10.1038/s41593-018-0101-9). Nat. Neurosci. *21*, 1018.
Pradeepa, M.M., Grimes, G.R., Kumar, Y., Olley, G., Taylor, G.C.A., Schneider, R., and Bickmore, W.A. (2016). Histone H3 globular domain acetylation identifies a new class of enhancers. Nat. Genet. *48*, 681–686.
Quinlan, A.R., and Hall, I.M. (2010). BEDTools: A flexible suite of utilities for comparing genomic features. Bioinformatics *26*, 841–842.
Radzisheuskaya, A., Shliaha, P. V., Grinev, V. V., Shlyueva, D., Damhofer, H., Koche, R., Gorshkov, V., Kovalchuk, S., Zhan, Y., Rodriguez, K.L., et al. (2021). Complex-dependent histone acetyltransferase activity of KAT8 determines its role in transcription and cellular homeostasis. Mol. Cell *81*, 1749-1765.e8.
Rai, T.S., Cole, J.C., Nelson, D.M., Dikovskaya, D., Mcbryan, T., Faller, W., Tuyn, J. Van, Morrice, N., Hewitt, R.N., Manoharan, I., et al. (2014). HIRA orchestrates a dynamic chromatin landscape in senescence and is required for suppression of neoplasia. 2712–2725.
Ramírez, F., Dündar, F., Diehl, S., Grüning, B.A., and Manke, T. (2014). deepTools: a flexible platform for exploring deep-sequencing data. Nucleic Acids Res. *42*, W187-91.

Schertzer, M.D., Thulson, E., Braceros, K.C.A., Lee, D.M., Hinkle, E.R., Murphy, R.M., Kim, S.O., and Vitucci, E.V.A.C.M. (2019). A piggyBac-based toolkit for inducible genome editing in mammalian cells. 1047–1058.
Zhang, Y., Li, T., Preissl, S., Amaral, M.L., Grinstein, J.D., Farah, E.N., Destici, E., Qiu, Y., Hu, R., Lee, A.Y., et al. (2019). Transcriptionally active HERV-H retrotransposons demarcate topologically associating domains in human pluripotent stem cells. Nat. Genet. *51*, 1380–1388.
